# Adenosine deficiency facilitates CA1 synaptic hyperexcitability in the presymptomatic phase of a knock in mouse model of Alzheimer’s disease

**DOI:** 10.1101/2024.04.24.590882

**Authors:** Mattia Bonzanni, Alice Braga, Takashi Saito, Takaomi C. Saido, Giuseppina Tesco, Philip G. Haydon

**Affiliations:** Department of Neuroscience, Tufts University, Boston, MA, USA; Centre for Cardiovascular and 811 Metabolic Neuroscience, Department of Neuroscience, Physiology & Pharmacology, University College London, London, WC1E 6BT, UK; Department of Neurocognitive Science, Institute of Brain Science, Nagoya City University Graduate School of Medical Sciences, Nagoya, Aichi 467-8601, Japan; Laboratory for Proteolytic Neuroscience, RIKEN Center for Brain Science, 2-1 Hirosawa, Wako, Saitama 351-0198, Japan

**Author notes:** equally contributed (Mattia Bonzanni, Alice Braga). Corresponding authors and Lead Contacts.

**Keywords:** Adenosine, Hyperexcitability, Alzheimer’s Disease, Asymptomatic stage of AD, Ketogenic diet, Entorhinal cortex-hippocampus axis

## Abstract

The disease’s trajectory of Alzheimer’s disease (AD) is associated with and worsened by hippocampal hyperexcitability. Here we show that during the asymptomatic stage in a knock in mouse model of Alzheimer’s disease (APP^NL-G-F/NL-G-F^; APPKI), hippocampal hyperactivity occurs at the synaptic compartment, propagates to the soma and is manifesting at low frequencies of stimulation. We show that this aberrant excitability is associated with a deficient adenosine tone, an inhibitory neuromodulator, driven by reduced levels of CD39/73 enzymes, responsible for the extracellular ATP-to-adenosine conversion. Both pharmacologic (adenosine kinase inhibitor) and non-pharmacologic (ketogenic diet) restorations of the adenosine tone successfully normalize hippocampal neuronal activity. Our results demonstrated that neuronal hyperexcitability during the asymptomatic stage of a KI model of Alzheimer’s disease originated at the synaptic compartment and is associated with adenosine deficient tone. These results extend our comprehension of the hippocampal vulnerability associated with the asymptomatic stage of Alzheimer’s disease.

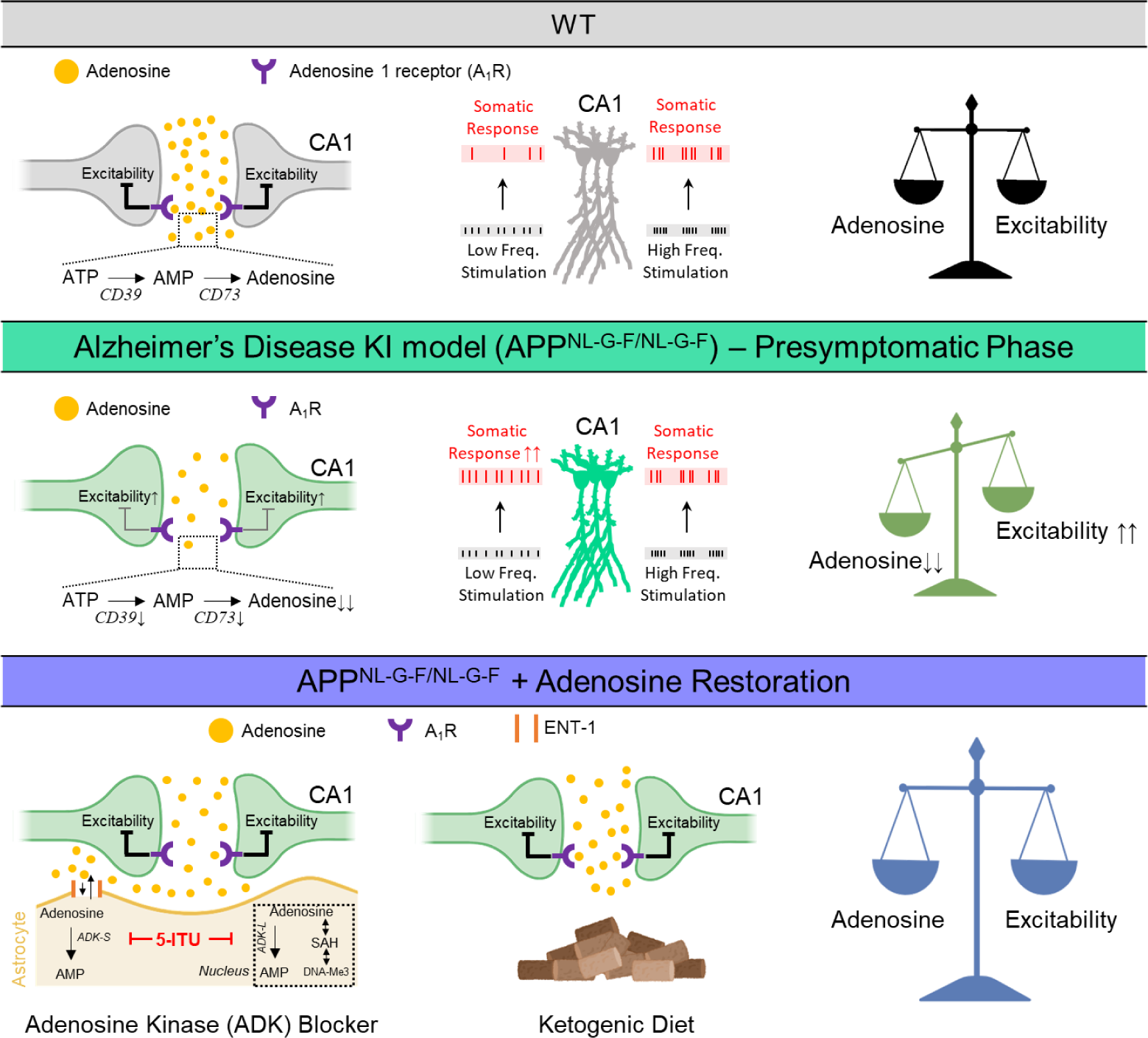

- Synaptic hyperexcitability spreads to the soma and manifests at low frequencies of stimulation
- Synaptic hyperexcitability is associated with reduced adenosine tone *in vitro* and *in vivo*
- Reduced adenosine tone is driven by reduced ATP-to-adenosine extracellular conversion
- Restoration of adenosine tone successfully normalizes neuronal excitability in the AD model

## Introduction

Alzheimer’s disease (AD) is a progressive neurodegenerative condition marked by the abnormal buildup of amyloid beta (Aβ) plaques, the formation of neurofibrillary tau tangles, and progressive neural degeneration^1^. Along with the hallmarks of the disease, neuronal hyperactivity has been reported in patients, animal models and in patient-derived cell cultures^2–12^. Previous studies^13^ have found that both sporadic and familial AD patients exhibit increased cortical-hippocampal activity, occurrence of seizures during the early phases of the prodromal stage^14–16^ and faster cognitive decline in patients that exhibit subclinical epileptiform activity^17,18^. Moreover, new evidence has linked Aβ-driven hyperactivity to subsequent aggregation and spreading of tau pathology^19^. Despite the evident comorbidity between neuronal hyperactivity and AD during the prodromic stage in patients, the origin of neuronal hyperexcitability and hippocampal vulnerability remain debated^20,21^. It is noteworthy that some reports have described hippocampal hyperactivity in asymptomatic offspring of autopsy-confirmed AD patients ^22,23^. Granted that such evidence is limited, it raises the hypothesis that hippocampal hyperexcitability may precede the prodromal stage also in humans. In fact, early-stage murine models of AD exhibit hippocampal hyperactivity^4,13,24,25^, which contributes to the deposition of amyloid beta and is promoted by amyloid beta toxicity^26^, calcium imbalance^27^, enhanced glutamate signaling^28^ and decreased GABA tone. Since it has been shown^29^ that breaking the vicious cycle between amyloid beta deposition and neuronal hyperexcitability ameliorated the disease’s trajectory, it is of relevance to unravel the players contributing to this aberrant activity at early stage of the disease. Since multiple mechanisms converge to control neuronal hyperactivity, additional contributors remain unexplored. Further investigation is needed to examine understudied factors contributing to the neuronal hyperexcitability during the asymptomatic (preclinical) phase of AD^1^, with the goal of understanding the origin of the functional hippocampal vulnerability during the earliest stages of AD.

Adenosine is a neuromodulator regulating both inhibitory and excitatory neuronal responses primarily through A_1_ and A_2A_ receptors^30^. Furthermore, adenosine influences the state of DNA methylation^31^. While dysregulated adenosine signaling, mainly acting through the A_2A_R axis, has been observed during the symptomatic stages of AD^32^, it is unclear whether adenosine signaling is affected during the asymptomatic stage of AD. Attempts to slow the progression of the disease by either chronically depressing^33^ or potentiating^34,35^ the adenosine signaling have produced contradictory results, highlighting the complexity of the issue. Adenosine functions as a “retaliatory metabolite,” serving as a feedback control mechanism in response to low energy states that may occur during aberrant neuronal activity. Given its anticonvulsant and antiepileptogenic properties^30^, along with our limited comprehension of the connection between adenosine signaling, hyperexcitability, and AD, we asked whether reduced adenosine signaling might participate in the hyperexcitable phenotype during the preclinical phase of AD.

In this work, we present evidence of a link between adenosine deficiency and hippocampal synaptic hyperactivity during the asymptomatic phase in a KI model of AD.

## Results

### CA1 synaptic activity is potentiated in the asymptomatic phase of the APPKI model

Synaptic alterations have been described as one of the earliest events in Alzheimer’s disease (AD) ^24^. Considering the vulnerability of the entorhinal cortex-hippocampus (EC-Hip) circuitry in both human and murine AD models^36^, we monitored the CA1 field, which serves as the converging site for two distinct pathways within the EC-Hip axis: the indirect pathway from CA3 to CA1 (Schaffer’s collateral (SC)) and the direct pathway from ECIII to CA1 (temporoammonic (TA) pathway). We tested the hypothesis that net synaptic transmission is potentiated in these two circuits in the APPKI model.

We monitored the CA1 *sr* (*stratum radiatum*) response by stimulating either the SC (Fig.1A-D) or TA terminals (Fig.1E-H) in the hippocampal slices from WT (black) and APPKI (green) mice at 8 weeks of age, an asymptomatic stage of the diseases with minimal to no plaque burden and absent tau pathology, gliosis, spine and cellular degeneration, and cognitive impairment^24^. We measured the evoked fEPSP (field excitatory post synaptic potential) and FV (fiber volley) responses as a proxy of postsynaptic activity and presynaptic recruitment, respectively. We then quantified net synaptic transmission of each slice as the slope of the FV-fEPSP relationship (Fig1B,F), where larger absolute values indicate an increased net synaptic transmission. As evident from the representative traces (insets) and the FV-fEPSP slope distributions, net synaptic transmission was potentiated in the APPKI model in both circuits (Fig.1B and Fig.1F, respectively); the FV response was not different between the genotypes, indicating that the same fraction of pre-synaptic fibers has been activated during stimulation (Figure S1). It follows that, given the same pre-synaptic activation (FV), the post-synaptic response (fEPSP) is potentiated in the APPKI model (Fig.1B,F, left).

**Fig. 1.**
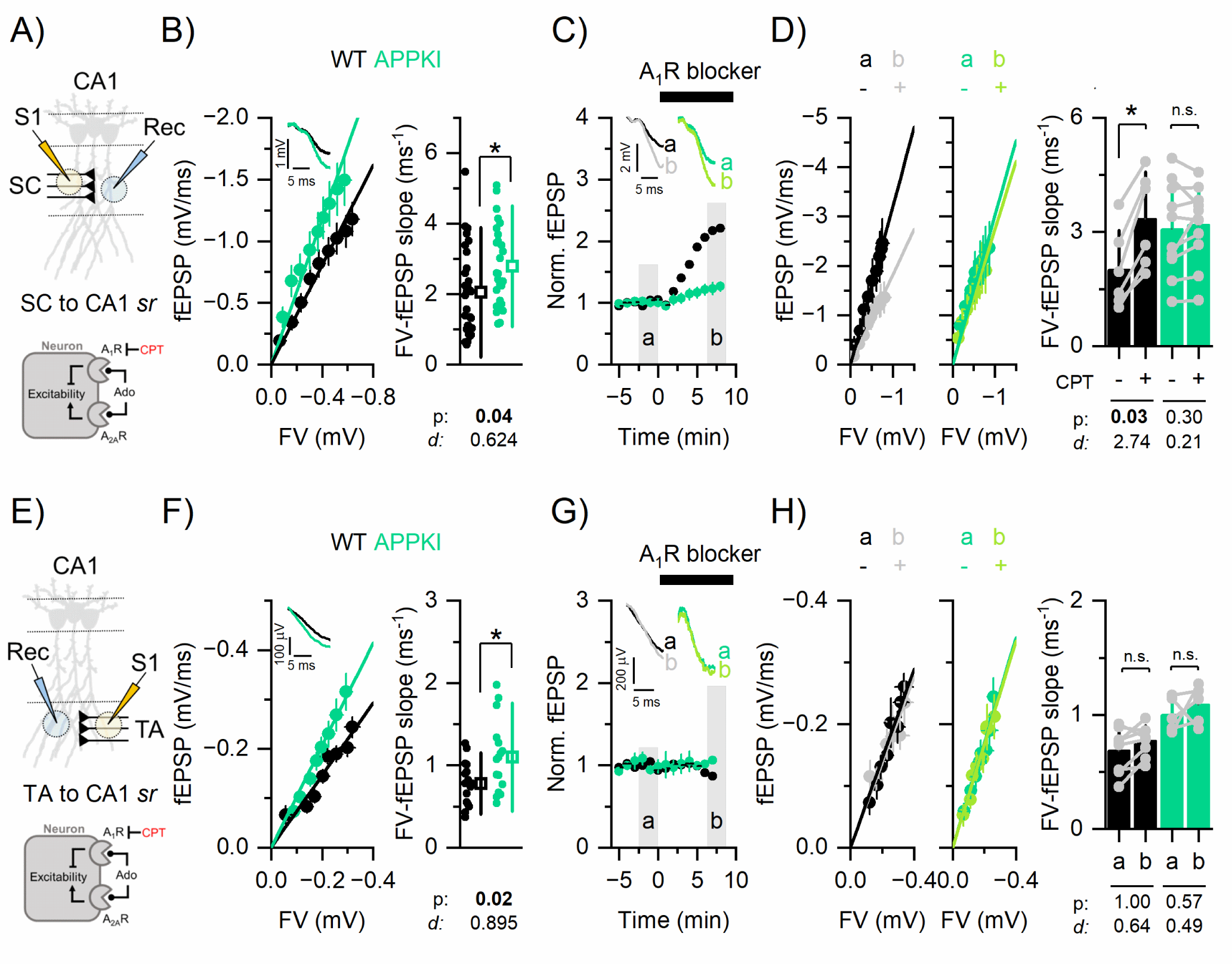
SC and TA synaptic signaling in the APPKI model is enhanced in an adenosine dependent and independent manner, respectively. **A,E**) Top: Scheme of experiment to monitor SC-to-CA1 and TA-to CA1 synaptic signaling (S1: stimulation; Rec: recording). Bottom: cartoon of adenosine signaling. **B,F**) Mean fEPSP-FV relationship (left) and individual slope data points (right); SC-to-CA1; WT: n=30/16, APPKI: n= 27/17; TA-to-CA1 - WT: n=18/11, APPKI: n= 16/9. Inset: representative fEPSP traces. **C,G**) Time trajectory of normalized fEPSP before (a) and after (b) the dosage of the A_1_R blocker, CPT (200 nM). **D,H**) Mean fEPSP-F.V. relationship (left) and individual paired slope data points (right) before (a) and after (b) dosage of the A_1_R blocker; SC-to-CA1; WT: n=12/3, APPKI: n= 20/6; TA-to-CA1 - WT: n=16/4, APPKI: n= 10/5. p: p-value; *d*: Cohen’s effect size. Mean±S.D. n=n. slices/n. mice. Statistic: generalized linear mixed effect models. Black: WT, Green: APPKI, Grey: WT+CPT, Light Green: APPKI+CPT.

These results demonstrated an increased net synaptic activity in both circuits in the absence of pre-synaptic fibers degeneration during the asymptomatic stage of AD.

### Enhanced synaptic transmission of SC synapses, but not TA synapses, is facilitated by adenosine deficiency in the APPKI model

Although various factors have been associated with neuronal hyperexcitability at different stages of AD^13^, the involvement of adenosine, a known endogenous anticonvulsant and antiepileptogenic neuromodulator, remains unexplored during the asymptomatic phase. We investigated whether a reduced adenosine 1 receptor (A_1_R)-dependent tone, known for its inhibitory role on neurons^30^, contributed to the enhanced synaptic transmission observed in the CA1 regions of the APPKI model.

To assess the A_1_R-dependent basal tone, we monitored the FV-fEPSP relationship before and after delivery of an A_1_R antagonist, CPT (8-cyclopentyl-1,3-dimethylxanthine, 200 nM). After computing the baseline FV-fEPSP relationship, we selected a stimulation intensity producing fEPSP with a slope that was 30-40% of the maximal and monitored the fEPSP before (a) and after (b) dosing the A_1_R blocker to confirm the steady state action of the drug (Fig.1C and Fig.1G). We then constructed the FV-fEPSP relationships in the presence of the blocker. In the SC to CA1 circuit, as evident from the time trajectory (Fig.1C) and further confirmed by the FV-fEPSP slope values (Fig.1D), net synaptic transmission was enhanced by the A_1_R antagonist in WT but not in APPKI. In the same circuit, we also confirmed that A_1_R receptors were functionally active by dosing an A_1_R agonist (Figure S2A-D) and found a similar decrease in net synaptic transmission in both models. Additionally, upon pharmacological inhibition of the ENT1 transporter (Figure S2E-H), which is primarily responsible for astrocytic adenosine uptake (its inhibition elevates the extracellular concentration of adenosine), we observed a reduction in net synaptic transmission in both models. This finding confirmed that the APPKI model retained the A_1_R-mediated inhibitory effects were still retained. Importantly, in the TA to CA1 circuit, both WT and APPKI were insensitive to the A_1_R antagonist (Fig.1G,H), suggesting that the TA to CA1 circuit was not under basal adenosine control.

These results revealed that a potential decrease in the baseline adenosine level, with intact A_1_R signaling, contributed to increased synaptic hyperexcitability at the SC terminals. Moreover, the lack of basal adenosine in the TA pathway emphasized the involvement of adenosine-independent factors in setting synaptic hyperactivity in the APP KI model.

### Heterosynaptic but not homosynaptic plasticity is reduced in the APPKI model

Considering the adenosine-mediated modification of synaptic activity at the SC synapses in the APPKI model, we investigated whether such alterations affected synaptic plasticity. We applied homosynaptic and heterosynaptic stimulation protocols to investigate the plasticity at the SC terminals.

We monitored the fEPSP time trajectory (Fig.2B) before (Fig.2B, a) and after (Fig.2B, b) high-frequency stimulation at the SC terminals (LTP protocol used as a proxy of long-term memory, Fig. 2A). The resulting synaptic potentiation was intact in both genotypes (Fig.2C). We then monitored the PPr (Paired Pulse ratio as a proxy of short-term memory; Fig.2E) finding a similar trend for the WT and APPKI models (Fig.2F). These results demonstrated that both long- and short-term cellular memory at the SC terminals were intact in the APPKI model.

**Fig. 2.**
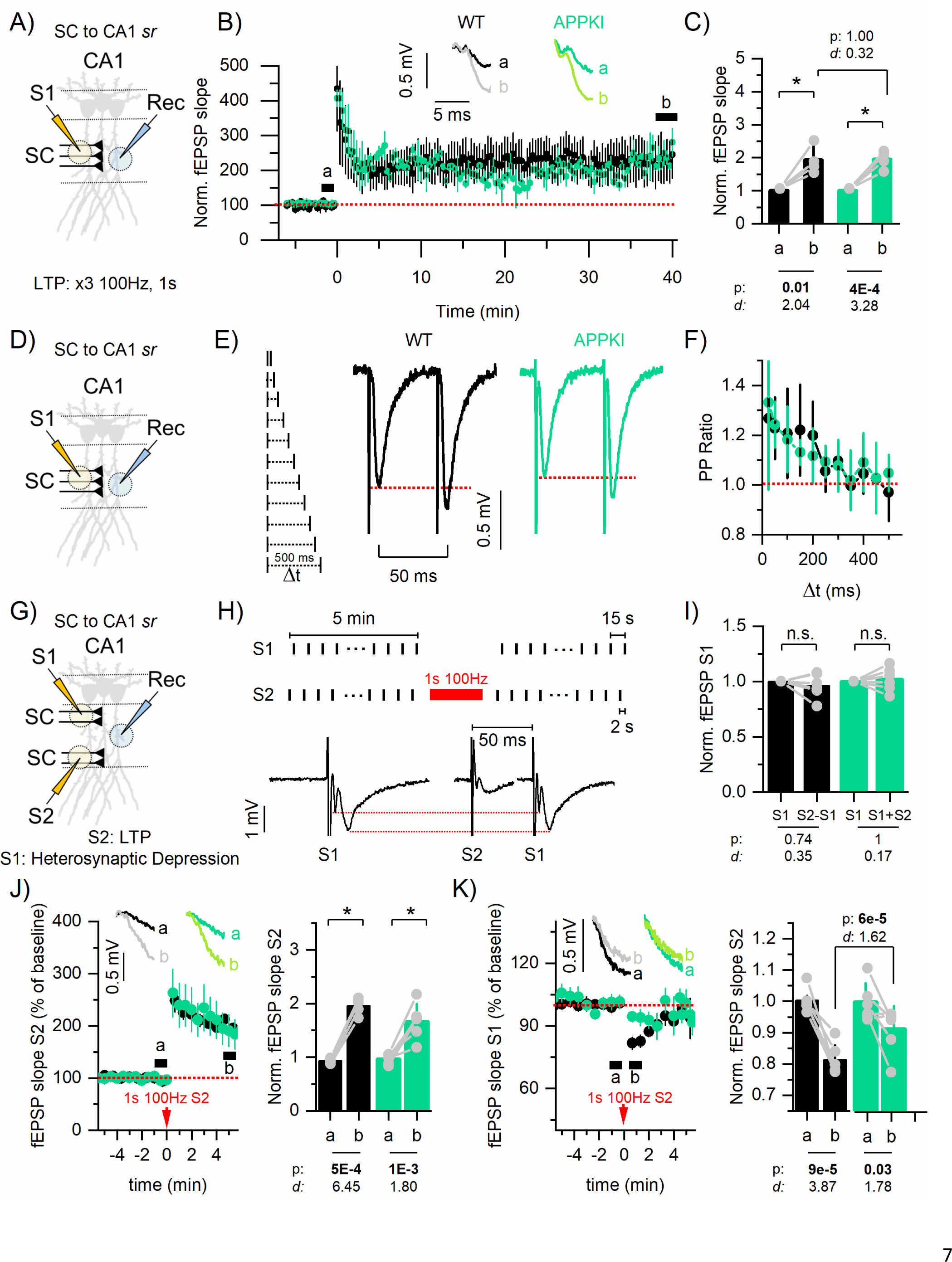
Heterosynaptic but not homosynaptic plasticity is affected at the SC terminals of the APPKI model. **A,D,G)** Scheme of experiment to monitor SC-to-CA1 and TA-to CA1 synaptic signaling (S1: stimulation fiber tract 1; S2: stimulation fiber tract 2; Rec: recording) **B)** Mean normalized fEPSP time trajectory before and after LTP protocol (at time=0 min; x3 100Hz stimulation, 1 second duration, 15 seconds inter-stimulus interval). Inset: representative fEPSP traces. **C)** Mean normalized fEPSP before (a) and after (b) LTP protocol and individual data points; WT: n=4/4, APPKI: n= 4/4. **E)** Paired pulse (PP) protocol with incremental inter-stimulation interval (Δt) (left) and representative traces at Δt=50ms (right). **F)** Mean PP Ratio as a function of Δt; WT: n=6/4, APPKI: n= 7/6. **H)** Heterosynaptic depression stimulation protocol (top) and representative traces (bottom) showing the independence of the two pathways S1 and S2; fEPSP elicited at S1 does not change when is preceded by S2 stimulation. **I)** Mean fEPSP of S1 alone and S2+S1 (Δt =50 ms) normalized on S1 alone; WT: n=6/6, APPKI: n= 6/6. **J)** Left: Mean normalized S2 fEPSP time trajectory before (a) and after (b) S2 1sec 100 Hz stimulation Inset: representative fEPSP traces. Right: Mean normalized S2 fEPSP before (a) and after (b) S2 stimulation and individual data points; WT: n=5/5, APPKI: n= 6/6. **K)** Left: Mean normalized S1 fEPSP time trajectory before (a) and after (b) S2 1sec 100 Hz stimulation. Inset: representative fEPSP traces. Right: Mean normalized S1 fEPSP before (a) and after (b) S2 stimulation and individual data points; WT: n=5/5, APPKI: n= 6/6. p: p-value; *d*: Cohen’s effect size. Mean±S.D. n=n. slices/n. mice. Statistic: generalized linear mixed effect models Black: WT, Green: APPKI.

Heterosynaptic depression at the SC terminals has been previously demonstrated to require an intact adenosine tone^37,38^. Therefore, we examined the hypothesis that the APPKI model lacked heterosynaptic depression as a result of the deficient adenosine tone. We stimulated two independent SC fibers (Fig.2G, S1 and S2); the S2 electrode served as site of a single high frequency burst (100 Hz, 1 s) resulting in transient potentiation of the synaptic transmission, while S1 served as site of heterosynaptic depression. To confirm pathway independence, we monitored the fEPSP amplitude at the S1 electrode either preceded or not by a 50-ms stimulation of the S2 electrode (Fig.2H). Our results (Fig.2I) indicated that the normalized fEPSP amplitude after S1 stimulation showed no significant difference, regardless of whether the S2 stimulation was performed or not, thus confirming pathway independence. In response to a 1s 100Hz stimulation at the S2 site (Fig.2J), synaptic activity at this site was similarly potentiated in both genotypes. At the S1 site synaptic depression was present in both genotypes, though significantly more prominent in the WT *vs* APPKI (Fig. 2K) model.

These results demonstrated that homosynaptic plasticity remains unaffected while heterosynaptic depression was reduced in the APP KI model. The evidence of a reduced heterosynaptic depression further supported the idea of a decreased adenosine tone since intact adenosine signaling is required for this form of plasticity.

### Enhanced CA1 somatic activity mirrors the synaptic potentiation in the APPKI model

Although WT and APPKI mice exhibited distinct synaptic responses, we asked whether somatic excitability was similarly enhanced. We first recorded the population spike (PS) at CA1 *pyr* (*stratum pyramidale*) in response to SC stimulation (Fig.3A,B). Net somatic activity was determined by monitoring the FV-PS amplitude relationship (Fig.3C). APPKI had increased net somatic activity (Fig.3C). The A_1_R antagonist CPT augmented PS in both genotypes although to different magnitudes (Fig.3D,E; see effect sizes). To ask whether the increased PS was a direct reflection of synaptic potentiation we also monitored the fEPSP-FV relationship (Fig.3F). When normalized by synaptic activity, the somatic output was not different between the two genotypes (Fig.3F). These results demonstrated that the enhanced synaptic activity at SC synapses is transmitted to the somatic compartment and that there is no further facilitation at the soma. These results suggested that the A_1_R-dependent hyperexcitable phenotype primarily occurs at the synaptic level.

**Fig. 3.**
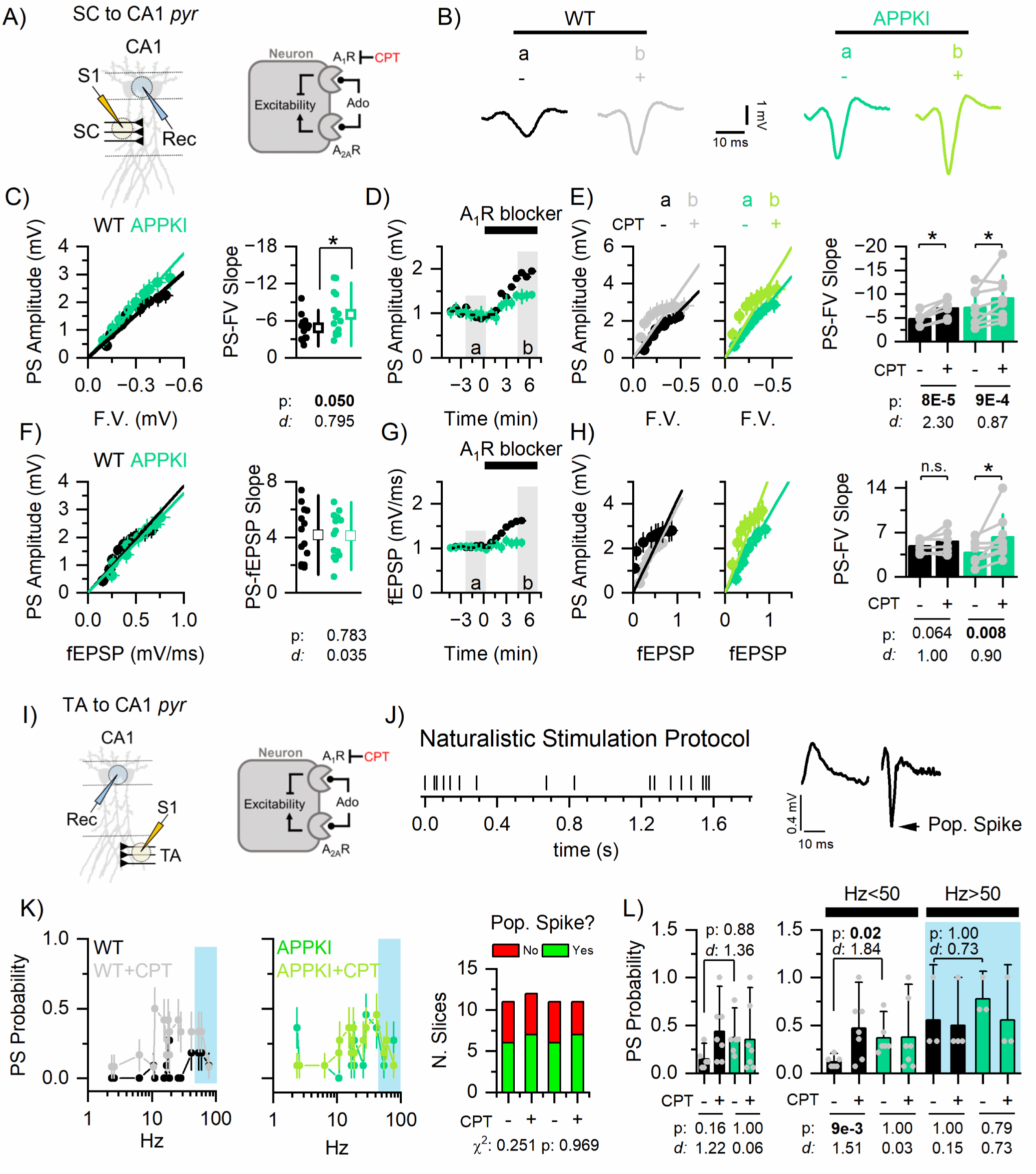
CA1 somatic hyperexcitability reflects synaptic potentiation in the APPKI model. **A,I)** Left: Scheme of experiment to monitor SC-to-CA1 and TA-to CA1 somatic signaling (S1: stimulation; Rec: recording). Right: cartoon of adenosine signaling. **B)** Representative population spike (PS) traces in the two genotypes before (a) and after (b) the dosage of the A_1_R blocker, CPT (200 nM). **C)** Mean PS Amplitude-F.V. relationship (left) and individual slope data points (right); WT: n=15/10; APPKI: n= 14/11. **D)** Time trajectory of PS amplitude before (a) and after (b) dosage of the A_1_R blocker. **E)** Mean PS Amplitude-F.V. relationship (left) and individual paired slope data points (right) before (a) and after (b) dosage of the A_1_R blocker; WT: n=6/5; APPKI: n= 8/5. **F)** Mean PS Amplitude-fEPSP relationship (left) and individual slope data points (right); WT: n=15/10; APPKI: n= 14/11. **G)** Time trajectory of fEPSP before (a) and after (b) dosage of the A_1_R blocker. **H)** Mean PS Amplitude-fEPSP relationship (left) and individual paired slope data points (right) before (a) and after (b) dosage of the A_1_R blocker; WT: n=6/5; APPKI: n= 8/5. **J)** Naturalistic stimulation protocol (left) and representative response without and with a population spike evoked (right). **K)** Mean probability of evoking a population spike (PS Probability) and fraction of slices in which at least 1 PS has been evoked; WT: n=6/4; WT+CPT: n=6/4; APPKI: n= 6/4; APPKI+CPT: n= 7/5. **L)** PS Probability of slices with at least 1 PS evoked without (left) or with a frequency-band segregation (High frequency: Hz>50; Low frequency: Hz<50). p: p-value; *d*: Cohen’s effect size. Mean±S.D. n=n. slices/n. mice. Statistic: generalized linear mixed effect models. Black: WT, Green: APPKI, Grey: WT+CPT, Light Green: APPKI+CPT.

To examine the somatic response in the TA pathway, we utilized a stimulus protocol that more faithfully resembles those found *in vivo* (naturalistic stimulation; Fig.3I,J), as sparse stimulation fails to elicit population spikes^39^. In Fig.3K, we report the probability of generating a population spike (PS Prob.) as a function of the frequency of stimulation. We first confirmed that the capability to generate population spikes during a naturalistic stimulation protocol was similar across conditions (WT, WT+CPT, APPKI, APPKI+CPT; Fig.3K). While the probability to generate a population spike during the naturalistic protocol was not different among conditions (Fig.3L, left), differences emerged after separating the stimulation into two frequency bands, non-burst and burst (below and above 50 Hz, respectively). As summarized in Fig. 3L (right), the likelihood of generating a population spike was not affected by either genotype or A_1_R blocker above 50 Hz (blue area). However, below 50 Hz, the probability of generating a population spike was significantly higher in the APPKI compared to WT. This enhanced probability in APPKI was due to an absence of adenosine-dependent suppression as revealed by the lack of action of the antagonist CPT. These results demonstrated that the TA pathway no longer exhibited high-pass filter properties in response to a naturalistic stimulation in an adenosine-dependent manner.

These results demonstrated an adenosine-dependent increased somatic activity of CA1 neurons in response to signals from both SC and TA terminals and the loss of the homeostatic control in the APPKI model.

### Hippocampal adenosine deficiency detected *in vivo* in the APPKI model

The functional data suggested that the adenosine levels might be decreased in the hippocampal brain slices of APPKI mice. To confirm this observation, we employed microdialysis to measure the *in vivo* adenosine levels in the hippocampus.

We implanted a guide canula in WT and APPKI mice at 7 weeks of age, after one week of recovery and 12 hours before dialysate collection, the probe was implanted targeting CA1 *sr* field (Fig.5A). We collected samples at ZT23-1, at the end of the dark phase when adenosine levels are at their maximum^40^. We found that both absolute levels of adenosine and levels normalized on the % of the animal activity were reduced in the APPKI model (Fig.4B-C).

**Fig. 4.**
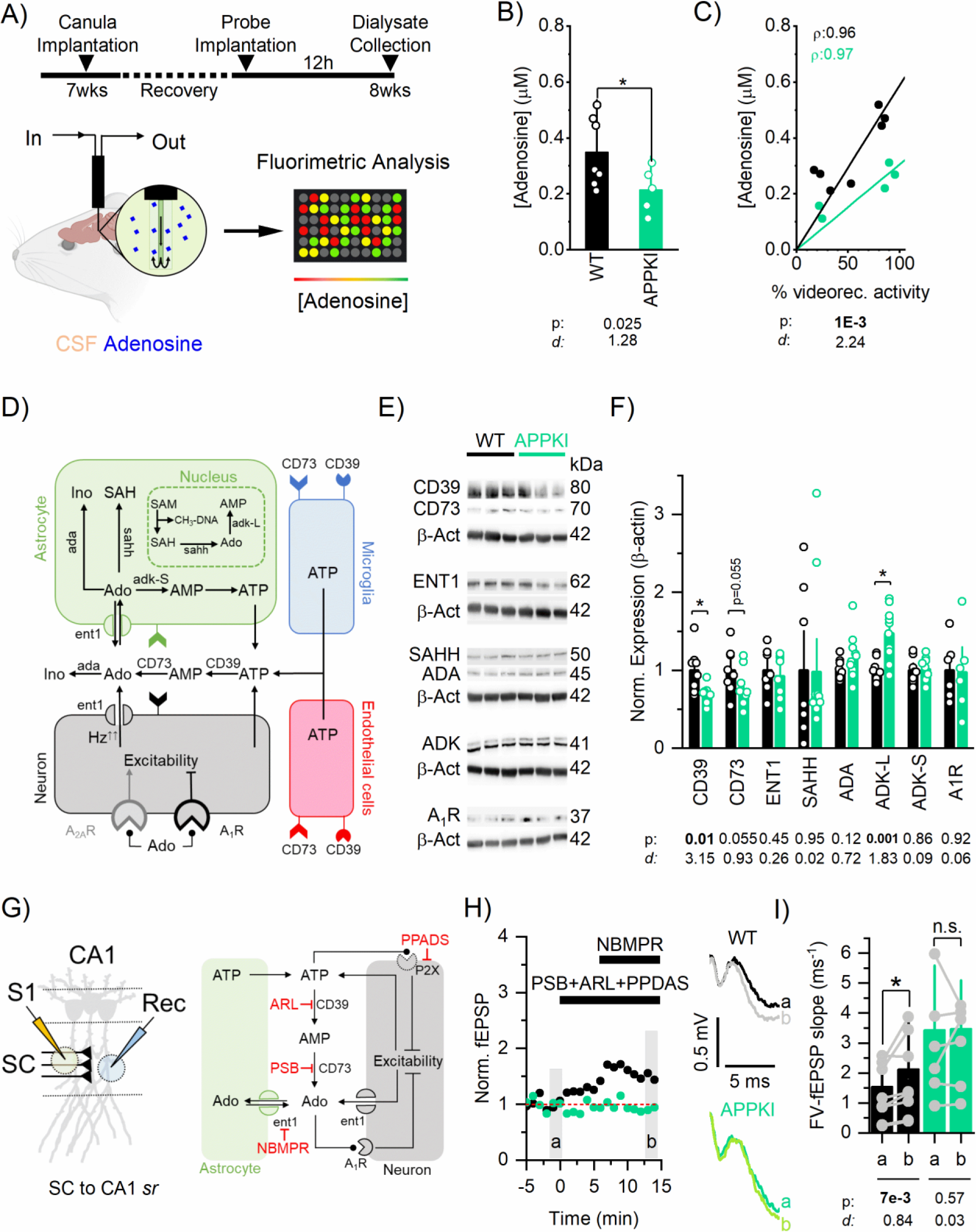
*In vivo* deficiency of adenosine is facilitated by reduced ATP-to-adenosine converting enzymes. **A)** Microdialysis and quantification protocol. **B)** Mean adenosine levels **C)** and relationship between adenosine concentration and percentage of activity during dialysate collection scored by video monitoring the mice (WT: n=7, APPKI: n= 5.). **D)** Cartoon of the main metabolic reactions and enzymes of adenosine pathway. **E)** Representative western blots and **F)** mean normalized expression and single data points. (CD39: WT: n=8, APPKI: n= 7; CD73: WT: n=8, APPKI: n= 7; ENT1: WT: n=7, APPKI: n= 8; SAHH: WT: n=8, APPKI: n= 11; ADA: n=9, APPKI: n= 12; ADK: WT: n=8, APPKI: n= 10; A1R: WT: n=6, APPKI: n= 6). **G)** Left: Scheme of experiment to monitor SC-to-CA1 synaptic signaling (S1: stimulation; Rec: recording). Right: cartoon of adenosine signaling and inhibitor targets (cocktail: ARL 67156 trisodium salt: 50 µM; PSB 12379: 50 nM; PPADS tetrasodium salt: 50 µM; NBMPR: 300 nM). **H)** Left: Time trajectory of normalized fEPSP before (a) and after (b) the dosage of the drug cocktail. Right: Representative traces before (a) and after (b) the dosage of the drug cocktail. **I)** Mean FV-fEPSP slopes before (a) and after (b) drug cocktail and individual data points; WT: n=8/8, APPKI: n= 6/6.p: p-value; *d*: Cohen’s effect size. Mean±S.D. n=n. slices/n. mice. Statistic: generalized linear mixed effect models. Black: WT, Green: APPKI. Grey: WT+cocktail, Light Green: APPKI+cocktail.

**Fig. 5.**
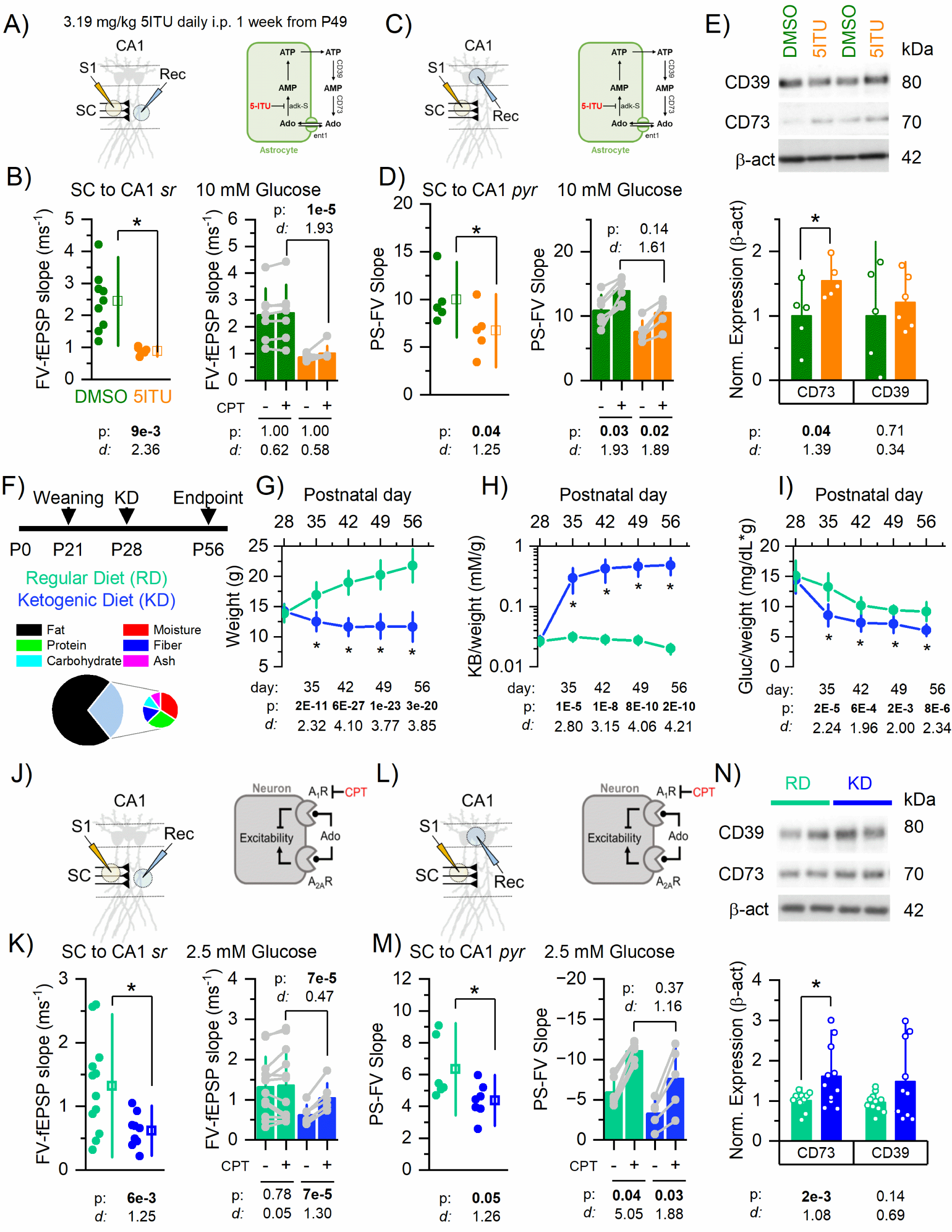
Normalization of neuronal hyperexcitability through pharmacological and ketogenic diet approaches. **A,C,J,L)** Left: Scheme of experiment to monitor SC-to-CA1 synaptic and somatic signaling (S1: stimulation; Rec: recording). Right: cartoon of adenosine signaling. **B)** FV-fEPSP data points (left) and individual paired slope data points (right) before (a) and after (b) dosage of the A_1_R blocker; DMSO: n= 9/4, 5-ITU: n= 7/6. **D)** PS-FV data points (left) and individual paired slope data points (right) before (a) and after (b) dosage of the A_1_R blocker; DMSO: n= 7/4, 5-ITU: n= 7/6. **E)** Top: Representative western blots. Bottom: Mean normalized expression and single data points (CD73: DMSO: n= 5, 5-ITU:n=5; CD39: DMSO: n= 5, 5-ITU:n=6). **F)** Regular diet (RD, green) and Ketogenic diet (KD. blue) regime and KD composition. At P28, KD group received *ad libitum* KD for a month (P56). Time trajectories of **G)** weight, **H)** ketone bodies/weight and **I)** glucose/weight for RD and KD groups. **K)** FV-fEPSP data points (left) and individual paired slope data points (right) before (a) and after (b) dosage of the A_1_R blocker; RD: n= 12/12, KD: n= 9/9. **M)** PS-FV data points (left) and individual paired slope data points (right) before (a) and after (b) dosage of the A_1_R blocker; RD: n= 6/6, KD: n= 7/7. **N)** Top: Representative western blots. Bottom: Mean normalized expression and single data points (CD73: RD: n= 11, KD: n=11; CD39: RD: n= 12, KD: n=10). p-value; *d*: Cohen’s effect size. Mean±S.D. n=n. slices/n. mice. Statistic: generalized linear mixed effect models. Dark Green: APPKI+DMSO, Orange: APPKI+5-ITU, Light Green: APPKI-RD, Blue: APPKI-KD.

These results supported our indirect adenosine assays that used synaptic and pharmacological approaches and demonstrate that the adenosine levels were reduced *in vivo* in the APPKI model.

### CD39/73 ATP-to-Adenosine machinery is impaired in the APP KI model

Considering the *in vivo* and *in vitro* evidence of decreased adenosine tone, our objective was to explore the underlying molecular factors contributing to this reduction. We thus analyzed the levels of enzymes involved in the synthesis, degradation, transport and signaling pathways of adenosine (Fig. 4D).

As summarized in Fig. 4E, we found that CD39, the rate-limiting enzyme of the two-step extracellular conversion reaction of ATP into adenosine, was significantly downregulated while ADK-L, the nuclear isoform responsible for the adenosine to AMP conversion, was significantly upregulated in the APPKI model. CD73, responsible for the second part of the ATP-to-adenosine reaction, was downregulated, albeit not significantly (p=0.055). Other enzymes involved in the degradation (SAHH, ADA and ADK-S), transport (ENT1) and signaling (A_1_R) were not differentially expressed in the two models. We then tested the capacity of the CD39/73 axis to generate adenosine under basal stimulation. We monitored the synaptic response at the SC terminals while simultaneously blocking CD39,CD73 and P2X receptors (to prevent ATP-mediated reduction of synaptic response through P2X receptors), followed by delivery of an ENT1 blocker (to prevent adenosine efflux through the equilibrative transporter after blocking extracellular adenosine production), as summarized in Fig. 4G. In the WT model, this cocktail increased net synaptic transmission by reducing ATP-mediated adenosine production. The antagonists were unable to change net synaptic transmission in the APPKI model, demonstrating the inefficiency of the CD39/73 axis to produce basal adenosine from ATP in this genotype (Fig.4.H,I).

These results demonstrated that the enzymatic machinery (CD39/73 axis) responsible for the extracellular conversion of ATP to adenosine was impaired while ADK-L level was upregulated in the APPKI model.

### ADK inhibition normalizes neuronal hyperexcitability in the APPKI model

Given the deficient adenosine tone and increased expression of ADK-L in the APPKI model, we investigated whether the transient administration of 5-iodotubercidin (5-ITU)^41^, a non-selective ADK inhibitor, could alleviate the hyperexcitable phenotype and restore the adenosine tone in the APPKI model. This strategy aimed to boost extracellular adenosine levels by inhibiting ADK-S and promote a shift in DNA methylation by inhibiting ADK-L. We either administered the drug (5-ITU) or the vehicle (DMSO) intraperitoneally for seven days daily starting at 7-weeks of age in the APPKI model. After a week of 5-ITU treatment, there was a significant reduction in both synaptic and somatic activity (Figure 5B-D). However, the treatment did not restore the adenosine tone synaptically (Fig.5B, left), likely attributed to the washout effect of the drug and its absence during brain slice experiments. In fact, experiments on untreated APPKI brain slices (see Figure S3) perfused acutely with 5-ITU confirmed the ability of 5-ITU to decrease excitability acting through A_1_R. Levels of CD73, as opposed to CD39, were increased in response to the *in vivo* administration of 5-ITU (Fig. 5E).

These results demonstrated that ADK inhibition during the asymptomatic phase produces long-lasting normalizing effects on neuronal excitability in the APPKI model.

### Ketogenic diet normalizes neuronal hyperexcitability and restore adenosine tone in the APPKI model

In addition to its well-recognized recognized positive impact on several AD-related markers ^42,43^, the ketogenic diet (KD) has demonstrated its ability to provide anticonvulsant and antiepileptogenic effects by acting through both adenosine-dependent and adenosine-independent pathways^44^. We therefore tested the hypothesis that ketogenic diet could normalize adenosine tone and neuronal hyperexcitability in the APP KI model.

One month of KD regime started at 4-weeks of age (Fig.5F) led to decreased weight (Fig.5G), increased circulating ketone bodies (Fig.5H; normalized on weight) and decreased circulating glucose (Fig.5I; normalized on weight) compared to age-matched mice fed with a regular diet (RD). The KD regime significantly reduced SC-to CA1 synaptic hyperexcitability (Fig.5K,M) and restored adenosine tone (Fig.5K) in the APPKI model. Similarly, KD reduced the somatic hyperexcitability while leaving intact the somatic adenosine response (Fig.5M). CD73 levels were restored by the KD regime, while CD39 levels were only restored in a sex-specific manner (Table S1).

These results demonstrated the efficacy of the ketogenic diet to restore adenosine tone and normalize neuronal excitability in the APPKI model.

## Discussion

The present research provides evidence of an association between hippocampal synaptic hyperactivity and adenosine deficiency during the asymptomatic phase in a KI murine model of Alzheimer’s disease (AD). Reversing aberrant excitatory activity ameliorates the AD trajectory^29^; it is thus relevant to study at which stage and which factors contribute to different forms of neuronal hyperexcitability during the AD trajectory. Here we focused on the asymptomatic phase (8 weeks) of the disease in an APPKI model aiming to elucidate novel factors contributing to the hippocampal vulnerability in AD.

In the hippocampus of APPKI mice, we found a potentiated synaptic basal response (SC and TA pathways) in line with previous works^24^. The somatic response scaled with the synaptic activity in both genotypes (Fig.3F), indicating that the enhanced synaptic response in the APPKI spread to the somatic compartment (Fig.3). Accordingly, previous works did not find intrinsic CA1 changes in firing, resting potential and input resistance^24^. This corroborated the findings that the synapse is an early locus of functional susceptibility. The functional distinction between genotypes became prominent primarily during low-frequency stimulations (Fig.1,3) while the response to high-frequency stimulations remained largely intact (Fig.2, 3L), as exemplified by the loss of high-pass filter properties in the TA pathway in APPKI CA1 neurons (Fig. 3L). Our findings align with the lack of hippocampal-related behaviors in aged-matched APPKI mice, as hippocampal computations are usually associated with sparse high-frequency neuronal activities. To further support this thesis, we also tested a stimulation protocol that engages both TA and SC terminals and mimics the temporal patterns and rhythmic nature (theta stimulation) of *in vivo* synaptic inputs to CA1 cells as mice traverse their place fields^45^ (Figure S4). We monitored two phenomena: phase precession and frequency adaptation. We confirmed that hippocampal theta-phase precession, a behavior involved in spatiotemporal coding and generated within the CA1 field *in vivo* ^46^, was observable *in vitro* and we found that was consistent across genotypes, whereas frequency adaptation appeared to be impaired in the APPKI genotype. This suggests that while the behavioral phenomenon remains intact in the APPKI model, sustaining it might demand higher energy expenditure. In general, even though there are no observable behavioral manifestations at this stage, damage is already occurring at the circuit level. The synaptic hyperactivity reported here and in other studies^24^is suggestive of an increased circuit noise. Under low-intensity and low-frequency presynaptic stimulations, the APPKI CA1 neurons may exhibit a larger and/or more frequent synaptic response compared to WT. This might indicate a compromised ability to filter incoming noisy synaptic communication, while still maintaining intact circuit-level functions associated with high-frequency activities. This increase in system noise level is compatible with the cascading network failure hypothesis^47^, which suggests that even in the absence of observable circuit-level phenotypes, an overloaded system becomes more vulnerable and prone to failure. If noisy signals aren’t effectively filtered out, this could worsen an ongoing energy crisis that has already been associated with the initial phases of Alzheimer’s disease development^48^.

Considering the A_1_R-mediated inhibitory action of adenosine on hippocampal neuronal activity (presynaptically and postsynaptically) ^30^, we showed that adenosine levels are reduced *in* vivo (Fig.5B-C), and that reduced adenosine tone enable hippocampal hyperexcitability. Our findings differed from those seen in later stages of the disease, where inhibiting the activity of adenosine receptors (such as with caffeine) improved synaptic plasticity and behavioral defects^33^. Previous studies showed increased levels of A_1_R in advanced disease’s stages and different models, including 5XFAD^49^, DeltaK280^50^, 3xTg^51^ and MAPT P301L^51^mice. However, during the asymptomatic stage, we observed intact functional A_1_R activation (Figure S2A) and A_1_R protein levels (Fig.4E) in the APPKI model. Interestingly, the somatic (Fig. 3D) but not the synaptic A_1_R-mediated response was intact, further implying pathology localized at the synaptic level. Aberrant A_2A_R-mediated signaling contributed to hippocampal deficits in several AD murine models^52–54^. The perfusion of the A_2A_R blocker in brain slices of the APPKI model (Figure S6) was ineffective in changing the net synaptic transmission, leading us to exclude the involvement of a potentiated A_2A_R-mediated signaling in the hyperexcitability phenotype.

Despite the TA pathway being unresponsive to the A_1_R blocker in both WT and APPKI models while under basal stimulation (0.033Hz), WT but not APPKI mice evoked adenosine-dependent responses while applying a naturalistic stimulation (Fig.3O). The synaptic amplitude recorded in the *pyr* layer was significantly larger in the APPKI *vs* WT and WT+CPT *vs* WT conditions, but not in the APPKI *vs* APPKI+CPT comparison (Figure S5), further supporting an impaired capacity to filter incoming low-frequency signals driven by the adenosine deficiency. The absence of a basal A_1_R-mediated signaling may be explained by reduced expression of A_1_R in the *slm* compared to the *sr* layer of the CA1 field in WT mice^55,56^. While we do not present a mechanistic explanation, we speculate that the adenosine-dependent potentiation of the somatic response could be the result of decreased filtering of the TA signal due a A_1_R-mediated change of the axonal resistance in the *sr* field. Alternatively, A_1_R-mediated effects may arise from adenosine accumulation in the *slm* layer.

Reduced adenosine tone was driven by reduced levels and activity of CD39/73 enzymes responsible for ATP-to-adenosine conversion, with reports indicating an increase in CD39/73 levels in more advanced stages of the disease^35^. Recent research emphasized the direct role of the microglial CD39/73 axis in facilitating the ATP-to-adenosine conversion and depressing neuronal excitability^57^. Notably, the levels of CD39 appear to positively correlate with the levels of P2Y12R, a homeostatic microglial marker^58,59^. Therefore, it would be important to examine the extent to which microglial activation (and concomitant loss of P2Y12R signature) contributes to the hyperexcitability phenotype observed in APPKI mice. In addition, we found ADK-L, the nuclear isoform of adenosine kinase involved in the DNA methylation process, upregulated in the APPKI model. Of notice, ADK-L is predominantly expressed in astrocytes in the adult mouse^60^, suggesting that hippocampal astrocytes in the APPKI model may exhibit DNA hypermethylation^61^. While the possibility of neuronal re-expression cannot be ruled out, our findings pave the way for new experimental questions investigating whether astrocytic DNA hypermethylation contributes to neuronal hyperactivity in Alzheimer’s disease, akin to epilepsy^61^.

The potentiation of adenosine tone has proved advantageous to normalize aberrant neuronal activity^44^.We found that a transient pharmacological restoration of the adenosine tone normalized neuronal excitability, even in the presence of other mechanisms driving the aberrant activity. We pharmacologically inhibited (5-ITU)^41^ the activity of ADK (adenosine deaminase kinase), while previous manipulations modulated ENT1 activity^34,35^. The administration of the 5-ITU treatment was performed intraperitoneally, making it difficult to rule out potential systemic effects. By selecting an ADK inhibitor we leveraged adenosine receptor-dependent and independent effects. In fact, blocking ADK-S (cytosolic isoform) activity rapidly elevates extracellular adenosine levels^31^, wheatear blocking ADK-L (nuclear isoform) activity decrease DNA methylation^41^. As the normalization of neuronal activity persisted beyond the duration of the acute drug’s effects (Figure S3), this suggested potential epigenetic effects likely involving inhibiting ADK-L and shifting astrocytic DNA methylation state.

Given the diverse array of metabolic effects associated with systemically and pharmacologically enhancing adenosine tone, it might not be prudent to employ this strategy as a potential therapeutic intervention in humans. We showed that adenosine levels can be restored, and excitability normalized using a non-drug treatment, the ketogenic diet, widening the array of neuroprotective effects of the diet in the AD context. Although the ketogenic diet has been shown to reduce neuroinflammation and plaque deposition in AD preclinical models ^42,62–65^, its effectiveness in restoring adenosine tone and normalize neuronal activity remained unexplored. The normalization of the adenosine tone is likely the results of well-known increasing of the ATP levels^42^ rather than a restoration of the CD39-73 axis. Since we fed the ketogenic diet to mice between one and two months of age, it is essential to noticing that this timeframe holds developmental significance, and any results obtained may be influenced by concurrent developmental changes.

In addition to the previously stated limitations, we used a murine model characterized by aggressive and rapid amyloid beta deposition. It is thus crucial to assess the generalizability of these results in models with a more physiological evolution of the disease. In the present study, we employed a WT murine model as a control of the APPKI. In addition to the familial mutations, the humanized version of the APP sequence (hAPP) further distinguished the two models. To address this, we replicated a subset of functional experiments in the hAPP model (Figure S7), which harbored the humanized APP sequence. The results demonstrated that in the hAPP model the adenosine tone was intact and the excitability physiological, ruling out any contribution of the humanized APP sequence as a driver of the observed differences.

The results of this study highlighted adenosine signaling as a vulnerability in early AD and, while this might suggest the augmentation of adenosine levels as a viable neuroprotective route of intervention, caution is warranted. While we found evidence of a neuroprotective role of adenosine augmentation during the asymptomatic stage of AD, other studies observed a beneficial effect by suppressing adenosine receptor signaling in later stages of the disease^33^. Rather than being conflicting, these results indicate that the neuroprotective effects of adenosine augmentation may transition to neurotoxicity depending on the stage of the disease. Moreover, identifying the optimal intervention window for adenosine augmentation currently presents a diagnostic challenge. Our results substantiate the need to further explore the causes and consequences of adenosine alterations in the early stages of disease in other murine models and human patients. In fact, comprehending the causes leading to altered adenosine tone (i.e., amyloid beta-driven cell state transitions, altered metabolism) might illuminate the earliest events of AD. At the same time, by investigating both adenosine receptor dependent and independent effects (in particular the astrocytic epigenetic effects) on neuronal excitability might provide more targeted strategies of intervention not only for AD, but also for other brain disorders.

In conclusion, we showed that adenosine deficiency compromise synaptic homeostasis contributing to CA1 neuronal hyperexcitability during the presymptomatic stage of an amyloid beta APPKI model.

## Acknowledgments

This work was supported by an award from the National Institutes of Health, 5R01AG061838 (to GT and PGH).

## Author contributions

Conceptualization, M.B, A.B. and P.G.H.; Methodology, M.B and A.B.; Software, M.B.; Investigation, M.B and A.B.; Formal Analysis, M.B.; Visualization, M.B and A.B.; Data Curation, M.B.; Writing – Original Draft, M.B, A.B. and P.G.H.; Writing – Review & Editing, M.B, A.B., T.C.S., T.S, G.T. and P.G.H.; Funding Acquisition, G.T. and P.G.H.; Resources, T.C.S., T.S and P.G.H.; Supervision, M.B and P.G.H.; Project Administration, M.B. and P.G.H.

## Declaration of interests

Nothing to declare.

## Lead contact

Further information and requests for resources and reagents should be directed to and will be fulfilled by the lead contacts, Mattia Bonzanni or Philip G. Haydon.

## Materials availability

This study did not generate new unique reagents.

## Data and code availability

All data points used in the figures, statistical models and raw western blot images are available at https://zenodo.org/records/10712113 (DOI: 10.5281/zenodo.10712113).

## Supplemental Information

Document S1. Figures S1-S7 and Table S1.

## STAR Methods

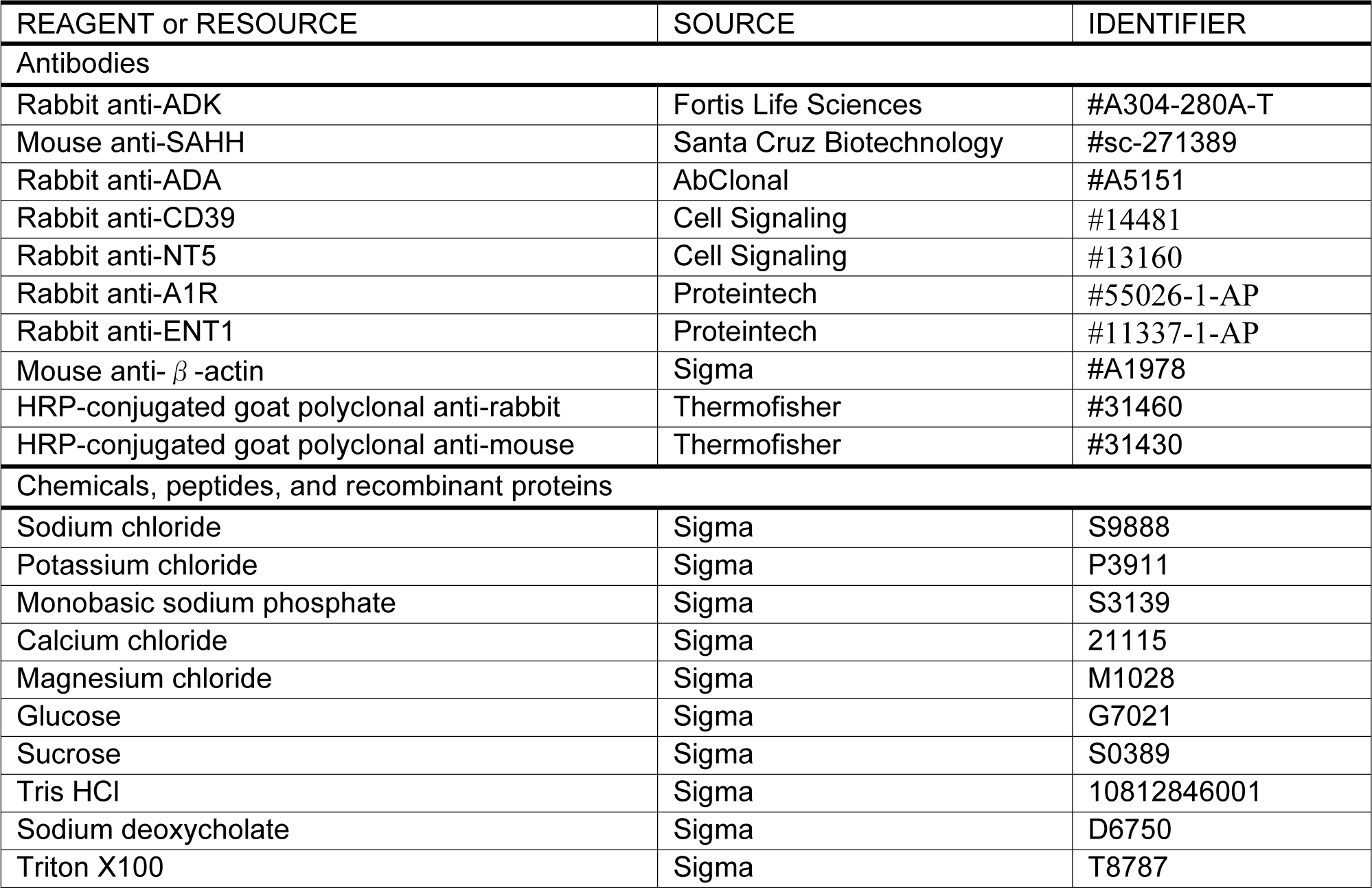
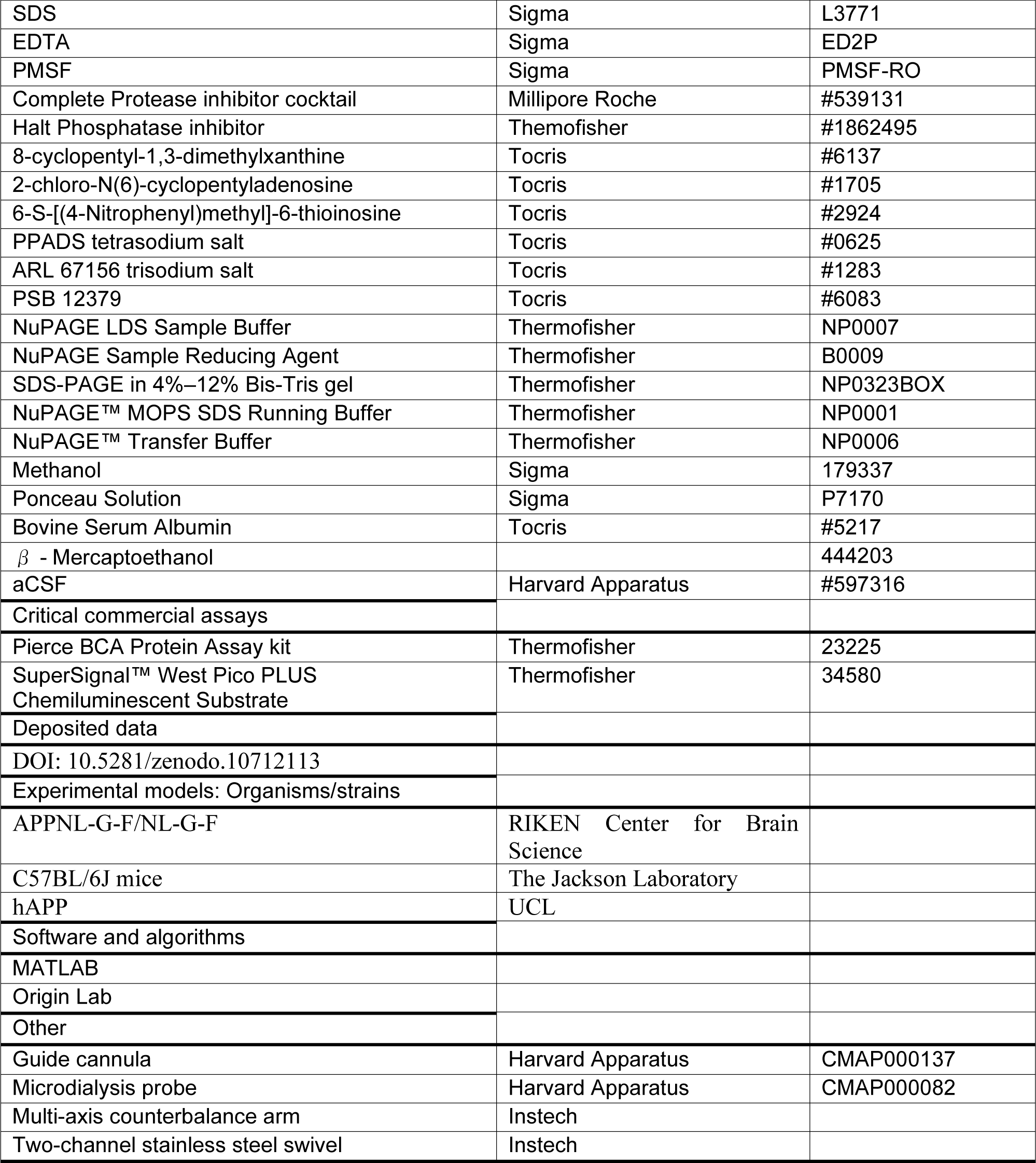

### Experimental Model

#### Animal

Homozygous APPKI mice carrying the humanized *App* gene with the Arctic, Swedish and Beyreuther/Iberian mutations (APP^NL-G-F/NL-G-F66^) were supplied by the RIKEN Center for Brain Science. Wild type (WT) C57BL/6J mice were purchased through Jackson Laboratory (Cat# 000664) and used as controls for the APPKI. To control for the humanized version of the *App* gene, a subset of experiments has been replicated using homozygous hAPP mice carrying the humanized *App* gene, which were supplied by UCL (Bart De Strooper). Adult mice were analyzed between 8 and 9 weeks of age (days: 57.10±2.80; mean±s.d.). This time point corresponded to minimal to no presence of amyloid burden, as well as the absence of tau pathology, gliosis, spine and cellular degeneration, and cognitive impairment in APPKI mice, as previously reported ^24,66^. Mice were bred and housed on a 12/12 light/dark cycle. We used the Zeitgeber time (ZT) scale that sets the origin of the 24h period (ZT0) to the onset of the light-phase (at ZT12 starts the dark phase), allowing comparison among studies independently of the actual clock-time settings of animal facilities. Mice were fed *ad libitum* with either regular chow (Regular diet; RD) or with Ketogenic diet (KD). KD is formulated as high fat, low carbohydrate diet in a paste form, with the ratio of fat to carbohydrate and protein approximately at 6:1 (F3666, Bio-Serv). KD was administered for a month starting at four weeks of age, with weight, ketone bodies and glucose monitored weekly. For the KD group, heavy loss of weight (more than 50%) and immobility within the first 3 days of the diet were used as exclusion criteria; for the APPKI genotype, about 10% of the mice were excluded within the first week of KD. In all genotypes, both sexes have been used and sex has been included as independent variable in all the analysis. Animals have been single housed only for the microdialysis experiment. For the diet and *in vivo* injection treatments, littermates have been randomly assigned to each group. All animal experiments were conducted in accordance with the guideline of the Animal Care and Use Committee of Tufts University.

### Method details

#### Tissue preparation

Hippocampal-entorhinal cortex (HEC) brain slices were prepared from 8 weeks-old male and female mice of the following genotypes: C57BL/6J, hAPP and hAPP^NL-G-F/NL-G-F^ (APP KI ^66^). Mice were anesthetized with isoflurane and rapidly decapitated. The brains were removed from the skull and placed in cold modified aCSF cutting solution with 10 mM of glucose (in mM: NaCl 120, KCl 3.2, NaH_2_PO_4_ 1, CaCl_2_ 1, MgCl_2_ 2, NaHCO_3_ 26, Glucose 10) or 2.5 mM glucose (in mM: NaCl 120, KCl 3.2, NaH_2_PO_4_ 1, CaCl_2_ 1, MgCl_2_ 2, NaHCO_3_ 26, Glucose 2.5, Sucrose 7.5). To prepare dorsal hippocampal-entorhinal cortex slices, which guarantees optimal preservation of the perforant path inputs ^67^, after removal of the olfactory bulb the brains were glued to an agar ramp (slope 10-12° with the anterior surface facing up the slope) and cut submerged under cold modified aCSF solution into 350 μm thick sections (Leica Vibratome). For electrophysiological experiments with 10 mM glucose, hemislices were then submerged into a storage container filled with aCSF (in mM: NaCl 120, KCl 3.2, NaH_2_PO_4_ 1, CaCl_2_ 2, MgCl_2_ 1, NaHCO_3_ 26, Glucose 10) at 35°C for 30 minutes and subsequently recovered at room temperature for at least 1 hour. For electrophysiological experiments with 2.5 mM glucose, hemislices were submerged into a storage container filled with aCSF (in mM: NaCl 120, KCl 3.2, NaH_2_PO_4_ 1, CaCl_2_ 2, MgCl_2_ 1, NaHCO_3_ 26, Glucose 2.5, Sucrose 7.5) at 35°C for 45 minutes and subsequently recovered at room temperature for at least 20 minutes. For western blot tissue preparation, after isolation the slices were moved into a vial containing RIPA (10 mM Tris HCl, 0.1 M, pH 7.2; 1% sodium deoxycholate; 1% Triton X-100; 1% sodium dodecyl sulfate [SDS]; 150 mM NaCl, 1.5 M; 1 mM EDTA, pH 8.0, 0.5 M (#AM9260G, Thermo Fisher Scientific); 1 mM phenylmethanesulfonyl fluoride (#93482, Sigma); Complete Protease Inhibitor cocktail (#539131, Millipore Roche); Halt Phosphatase Inhibitor (#1862495, Thermo Fisher Scientific)). All the solutions were constantly bubbled with 95%-5% O_2_-CO_2_ mix. All tissue collections have been performed between ZT-23 and ZT-1, time point at which adenosine concentrations are maximal.

#### Electrophysiology

Brain hemislices were transferred to a recording chamber mounted on a microscope (Olympus Microscope BX51) and superfused with aCSF saturated with 95%-5% O_2_-CO_2_. All field potential recordings were performed at 33°±1°C with an extracellular glass pipette (7-10 MΩ) filled with aCSF and stimulations were delivered using bipolar tungsten electrodes (FHC #30200). Glutamatergic/GABAergic signals and CA3-to-CA1 connections were left intact in all recordings.

#### Pharmacology

The A_1_R-mediated inhibitory tone was measured by perfusing the slices with the A_1_R antagonist 8-cyclopentyl-1,3-dimethylxanthine (CPT, 200 nM; Tocris Bioscience). Sensitivity to A_1_R activation was estimated using the A_1_R-specific agonist 2-chloro-N(6)-cyclopentyladenosine (CCPA, 10 nM; Tocris Bioscience). Passive production and accumulation of extracellular adenosine was measured indirectly by blocking the equilibrative nucleoside transporter 1 (ENT1) using the transport inhibitor 6-S-[(4-Nitrophenyl)methyl]-6-thioinosine (NBMPR, 100 nM; Tocris Bioscience). PSB 12379 (50 nM; Tocris Bioscience) has been used as CD73 inhibitor. ARL 67156 trisodium salt (50 μM; Tocris Bioscience) has been used to inhibit CD39, since it is a non-selective NTPDase inhibitor. PPADS tetrasodium salt (50 μM; Tocris Bioscience) has been used as non-selective P2 antagonist to prevent the effect of ATP increase in response to the block of the ATP-to-adenosine conversion by PSB+ARL. 5-Iodotubercidin (5-ITU; Tocris Bioscience) has been used as adenosine kinase inhibitor.

#### Western Blot

Brain slices were prepared as described above. Total protein was extracted by resuspending the isolated tissue in radioimmunoprecipitation assay (RIPA) buffer and stored at -20°C until usage. Samples were then subjected to sonication (Soniprep 150, MSE). Samples were then spun at max vel for 10 min, and supernatants were collected for protein quantification. Protein quantification was performed using a Pierce BCA Protein Assay kit (#23227, Thermo Scientific). 20 μg of total protein were mixed with NuPAGE LDS Sample Buffer (#NP007, Life Technologies), NuPAGE Sample Reducing Agent (#NP0009, Life Technologies), and distilled water prior to being heated at 95°C for 5 min. Proteins were separated by SDS-PAGE in 4%–12% Bis-Tris gel (#NP0336, Thermofisher) using a Novex Bolt Mini Gel system and NuPAGE™ MOPS SDS Running Buffer (NP0001, Thermofisher) before being transferred onto Immobilon-P polyvinylidene fluoride membranes (#IPVH00010, 0.45 mm pore size; Millipore) in NuPAGE™ Transfer Buffer with methanol 20% (NP0006, Thermofisher) for 1 h 45 min at 4°C using a Novex Bolt Mini Blot Module. SeeBlue Plus2 standard (#LC5925, Life Technologies) was used to estimate protein sizes, and transfer was confirmed by Ponceau S (#BP103-10, Fisher Biotech) staining. Immunoblot was obtained by first blocking membranes for 1 h at room temperature with a solution of 0.1% Tween 20 and 5% Bovine Serum Albumin (BSA) (#5217, Tocris) in 1× PBS and then incubated with primary antibodies overnight at 4°C (rabbit anti-ADK 1:2000, #A304-280A-T, Fortis Life Sciences; mouse anti-SAHH 1:500, #sc-271389, Santa Cruz Biotechnology; rabbit anti-ADA 1:1000, #A5151, AbClonal; rabbit anti-CD39 1:1000, #14481, Cell Signaling; rabbit anti-NT5 1:1000, #13160, Cell Signaling; rabbit anti-A1R 1:1000, #55026-1-AP, Proteintech; rabbit anti-ENT1 1:500, #11337-1-AP, Proteintech). After washing with 1× PBS Containing 0.1% Tween 20, the membranes were incubated with the species-appropriate horseradish peroxidase-conjugated secondary antibodies (HRP-conjugated goat polyclonal anti-rabbit #31460, Thermofisher; or HRP-conjugated goat polyclonal anti-mouse #31430, Thermo Scientific) for 1 h at room temperature at a 1:20,000 dilution in blocking solution. Immunoreactivity was revealed using SuperSignal™ West Pico PLUS Chemiluminescent Substrate (#34577, Thermofisher) and imaged using a Fujifilm LAS 4000 Gel Imager system with ImageQuant LAS 4000 software (Fujifilm). If needed, antibodies were stripped from membranes prior to incubation with another primary antibody or prior to incubation with mouse anti-β-actin (1:1000, #A1978, Sigma) in a stripping solution (stripping solution 10%: 7.5% glycine, SDS 1%) with 0.7 β - mercaptoehtnaol (#63689, Sigma).

#### Microdialysis

7 weeks old male and female APPKI or WT mice were anesthetized with isoflurane, injected with buprenorphine (0.5mg/kg) and placed into a stereotaxic frame. One hole was drilled in the skull (hippocampus coordinates: AP=-2.5, ML=2 mm) and the guide cannula (#CMAP000137, Harvard Apparatus) was lowered into the hole to reach the coordinate DV =-1.7 mm upon insertion of the microdialysis probe (#CMAP000082, Harvard Apparatus). One anchor screw was inserted in the frontal area to ensure major stability of the system. Anchor screw and guide cannula were then secured to the skull with dental acrylic. 4 days after surgery, mice were moved into an insulated sound-proof chamber and placed into individual Plexiglas circle boxes (Pinnacle Technology, Lawrence, KS) containing water and food ad libitum. Mice were left 3 days for habituation prior to data collection and maintained on a 12:12 light/dark cycle. 17-18hours before the sample collection at the onset of the light phase (ZT23), the microdialysis probe was inserted into the guide cannula and connected to a two-channel stainless steel swivel (#375/D/22QM, Instech) attached to a multi-axis counterbalance arm (#MCLA/SMCLA, Instech) to allow free movements of the animal. ACSF (#597316, Harvard Apparatus) was perfused at a rate of 0.2µl/minute during the 17-18hours prior to sample collection and then switched to 1µl/minute at ZT23. Dialysis samples were stored at - 80°C until usage. During sampling, mice were video recorded (Sirenia software, Pinnacle Technology) and subsequently scored as active (mobile or awake) or as inactive (immobile and nested).

#### Ketone bodies and glucose blood levels

Blood samples were collected weekly to monitor ketone bodies and glucose levels with test strips (Abbott Diabetes Care). Blood was obtained via the tail snip method, which involved applying local disinfectant and anesthesia to the tail, followed by making a 1 mm incision with a scalpel blade, and stopping blood flow by gently dabbing the tail tip. Mice were observed for signs of infection over the subsequent three days.

#### Intraperitoneal injections

Mice received intraperitoneal (IP) injections of either DMSO or 5-ITU over a 7-day period, starting at 7 weeks of age, administered between ZT-9 and ZT-10. To minimize stress associated with handling, mice were acclimatized to the operator’s touch three times a week starting a week before injections. Following the injections, mice were observed for two hours. Those injected with 5-ITU exhibited reduced mobility, slower heart rate, and shallow breathing for approximately 4 hours post-injection. Criteria for exclusion included sustained immobility and weight loss within the initial two days, resulting in the exclusion of about 5% of mice from the 5-ITU group.

## Quantification and statistical analysis

### Electrophysiology

#### Synaptic basal response

field excitatory post synaptic potentials (fEPSP, a proxy of post-synaptic activity) from the CA1b *stratum radiatum* (*sr*) were recorded stimulating the Schaffer collaterals (SC) near the CA1a border at 0.067 Hz (every 15 seconds). fEPSP from the CA1 *stratum lacunosum moleculare* (*slm*) were recorded stimulating the medial entorhinal cortex layer 3 fibers (temporoammonic pathway; TA) in the CA1 *slm* at 0.067 Hz. Stimulation intensities were delivered in the range 20-100 μA with 10 μA of increment; each stimulation was repeated 4 times and the resulting traces averaged for analysis. Fiber volley (FV, a proxy of pre-synaptic recruitment) amplitude and fEPSP slope (measured in the first 10-30% of the trace after fiber volley to exclude contamination from somatic activation) were used to reconstruct the FV-fEPSP relation and the resulting slope was used as a proxy of net synaptic activity.

#### Homosynaptic plasticity

fEPSP recordings were taken from the CA1b *sr* by stimulating the Schaffer collaterals near the CA1a border at 0.067 Hz. Stimulation intensities were chosen to produce fEPSP with a slope that was 30-40% of those obtained with maximal stimulation for long-term potentiation (LTP) and paired-pulse ratio (PPr) protocols. The baseline recordings (stimulation at 0.067 Hz) were monitored for 15 minutes to ensure a stable fEPSP response. LTP was electrically induced at the Schaffer collaterals and recorded at CA1b *sr* by delivering three high-frequency stimulations (HFS, each consisting of 1 s duration 100 Hz train) at 0.033 Hz. fEPSPs were measured for 1 hour using stimulation of the fibers at 0.067 Hz. The fEPSP slope values were normalized by the average fEPSP values of the pre-stimulation. Analysis was conducted comparing the value of the fEPSP slope pre-stimulation (averaging the last five minutes before HFS) and after 1 hour from stimulation (averaging the last five minutes of the hour-long recording post-stimulation). PPr was examined at the CA1 *sr* by stimulating the Schaffer collaterals and monitoring the fEPSP amplitude of two consecutive stimulations with intervals in the range 25-500 ms. After normalizing the fEPSP slopes by the average of the pre-stimulus values, data were expressed as fold change (PPr Ratio) as a function of the interval (ms) between the two consecutive stimulations.

#### Heterosynaptic plasticity

heterosynaptic depression recordings were performed by placing two stimulating electrodes in two independent Schaffer collaterals (S1 and S2) and the recording pipette in the CA1b *sr*. Stimulation intensities were chosen to produce fEPSP with a slope that was 20-30% of those obtained with maximal stimulation for both electrodes. Pathway independence was confirmed by monitoring fEPSP amplitude after stimulating the two pathways (S2 preceding S1) at 50 ms interval. Baseline fEPSP slope was monitored for 10-15 minutes after reaching stable responses by stimulating S1 and S2 pathways (10 s interval) at 0.067 Hz. A 100 Hz 1 s stimulation was then delivered to S2. LTP and heterosynaptic depression were recorded from S2 and S1, respectively, at 0.067 Hz for 10 minutes. After normalizing the fEPSP slopes by the pre-stimulus values average, the trajectories were reconstructed. Analysis was conducted by averaging the minute before and after the 100 Hz stimulation in both pathways and normalizing the post-stimulation by the pre-stimulation values.

#### Basal somatic response

Population spikes from the CA1 *stratum pyramidale* (*pyr*) were recorded stimulating the Schaffer collaterals near the CA1a border at 0.067 Hz. Stimulation intensities were delivered in the range 20-100 μA with 10 μA of increment; each stimulation was repeated 4 times and the resulting traces averaged for analysis. FV, fEPSP slope and population spike amplitude (PS, a proxy of somatic activity) were monitored. FV (a proxy of pre-synaptic recruitment) amplitude and PS amplitude were used to reconstruct the FV-PS relation and the resulting slope was used as a proxy of net somatic activity. fEPSP slope and PS amplitude were used to reconstruct the fEPSP-PS relation and the resulting slope was used as a proxy of net synaptic transmission to the somatic compartment.

#### Naturalistic somatic response

fEPSP from the CA1 *pyr* were recorded stimulating the MEC3 fibers (TA pathway) using a naturalistic stimulation protocol (Inter-stimulation interval (s): 0, 0.0532, 0.0661, 0.1046, 0.1394, 0.1927, 0.2862, 0.6734, 0.8293, 1.249, 1.2734, 1.3633, 1.4220, 1.4771, 1.5413, 1.5596, 1.5762. Instantaneous frequency (Hz): 18.79, 77.86, 25.95, 28.68, 18.79, 10.69, 2.58, 6.41, 2.38, 41.92, 11.12, 17.03, 18.17, 15.57, 54.50, 60.56) adapted from Sun *et al*^39^. Stimulation intensities were chosen to produce fEPSP with an amplitude of 0.3-0.35 mV (0.32±0.11 mV). The probability of evoking a population spike (PS Probability) was quantified for each stimulus by visually identifying PS; the average probability of evoking a population spike was computed for each slice and compared among conditions. PS Probability was also plotted based on frequency profile and clustered in burst (Hz>50) *vs* non-burst (Hz<50) conditions and group comparisons evaluated.

#### Somatic heterosynaptic response

fEPSP from the CA1 *pyr* were recorded stimulating the TA and Schaffer collateral fibers with a theta protocol adapted from Milstein *et al*^45^. In detail, five theta cycles (150 ms each), bursts of five stimuli were delivered at 67 Hz (inter-stimulus interval of 15 ms) to both electrodes, with TA stimuli preceding Schaffer collateral stimuli by 20 ms. Stimulation intensities were chosen to produce fEPSP with an amplitude of 0.2-0.25 mV by stimulating the TA terminals and to produce fEPSP without a recognizable PS by stimulating the CA3; on occasions, the minimal stimulation could still elicit a small PS. PS was visually identified; the phase of the first evoked PS in the five theta cycles was evaluated (0.42 ms equivalent to 1°). The frequency of PS was calculated for each cycle and plotted against cycle number. Adaptation index 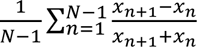 (where *N* is the total number of events and *x* is the chosen parameter) was used to quantify the phase and frequency adaptations over the five theta cycles.

### Western Blot

Densitometry measurements were performed using Fiji, with each protein band being normalized to their respective β-actin. For beautification purposes, WB bands in the main figures were cropped. Original blot images are available at https://zenodo.org/records/10712113 (DOI: 10.5281/zenodo.10712113).

### Microdialysis

Adenosine quantification has been conducted using the fluorometric Adenosine Assay Kit (Fluorometric) (Abcam; ab211094). Undiluted microdialysate has been used for the quantification.

### Statistic

Statistical analysis has been conducted using generalized linear mixed-effect models (glme) in MATLAB. glme offers a versatile approach for analyzing data deviating from a normal distribution, exhibits heteroskedasticity, and shows dependencies among observations (i.e., nested sampling: multiple brain slices from the same mouse)^68^. For each model, the distribution (Normal, Gamma, Inverse Gaussian) and fit method (REMPL, MPL) have been selected by choosing the model maximizing the relative likelihood (i.e., ℒ_*T*_ = *e*^0.5·(*AIC_min_*−*AIC_i_*)^, where *AIC_min_* is the minimum AIC across all models and *AIC_i_* is the AIC for the ith model) and visual analysis of the residual. By selecting the model with the largest ℒ_*T*_, we selected the model, among the tested models, that most strongly limits the information loss. For each panel, the number of slices and number of animals are reported in the caption, while p-values and effect sizes (*d* Coehn’s effect size reported as absolute value) are reported in the figure. For independent samples, the effect size has been computed as 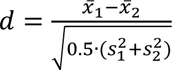, where 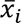 is the average of the i^th^ group and *s*_*i*_ is the standard deviation of the i^th^ group. By using the averaged variance at the denominator, we did not assume homogeneity of variance between groups. Hedge’s correction has not been applied. For paired samples, the effect size has been computed as 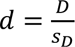, where *D* is the averaged difference between paired samples (*D* = 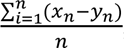, where *n* is the number of paired measures, *x* is group 1 and *y* is group 2) and *s*_*D*_ is the standard deviation of *D*. For each panel, the dependent and independent variables, and statistical details (F-Statistic, degrees of freedom, p-values) have been summarized in Table S1. For each panel, the model, dataset and modeling parameters are available at https://zenodo.org/records/10712113 (DOI: 10.5281/zenodo.10712113). Post-doc analysis has been performed using Bonferroni’s correction with alpha=0.05. No statistical measures were used to estimate sample size, since the effect size was unknown. Experiments were conducted by an investigator with knowledge of the animal genotype and treatment. Investigators were blinded during the analysis.

## Notes

https://zenodo.org/records/10712113

### Competing Interest Statement

The authors have declared no competing interest.

## References

1 van der Flier, W. M., de Vugt, M. E., Smets, E. M. A., Blom, M. & Teunissen, C. E. Towards a future where Alzheimer’s disease pathology is stopped before the onset of dementia. Nature aging 3, 494–505, doi:10.1038/s43587-023-00404-2 (2023).

2 Palop, J. J. et al. Aberrant excitatory neuronal activity and compensatory remodeling of inhibitory hippocampal circuits in mouse models of Alzheimer’s disease. Neuron 55, 697–711, doi:10.1016/j.neuron.2007.07.025 (2007).

3 Vossel, K. A. et al. Seizures and epileptiform activity in the early stages of Alzheimer disease. JAMA Neurol 70, 1158–1166, doi:10.1001/jamaneurol.2013.136 (2013).

4 Busche, M. A. & Konnerth, A. Neuronal hyperactivity--A key defect in Alzheimer’s disease? Bioessays 37, 624–632, doi:10.1002/bies.201500004 (2015).

5 Palop, J. J. & Mucke, L. Network abnormalities and interneuron dysfunction in Alzheimer disease. Nat Rev Neurosci 17, 777–792, doi:10.1038/nrn.2016.141 (2016).

6 Zott, B., Busche, M. A., Sperling, R. A. & Konnerth, A. What Happens with the Circuit in Alzheimer’s Disease in Mice and Humans? Annu Rev Neurosci 41, 277–297, doi:10.1146/annurev-neuro-080317-061725 (2018).

7 Ghatak, S. et al. Mechanisms of hyperexcitability in Alzheimer’s disease hiPSC-derived neurons and cerebral organoids vs isogenic controls. Elife 8, doi:10.7554/eLife.50333 (2019).

8 Zott, B. et al. A vicious cycle of beta amyloid-dependent neuronal hyperactivation. Science 365, 559–565, doi:10.1126/science.aay0198 (2019).

9 Babiloni, C. et al. What electrophysiology tells us about Alzheimer’s disease: a window into the synchronization and connectivity of brain neurons. Neurobiol Aging 85, 58–73, doi:10.1016/j.neurobiolaging.2019.09.008 (2020).

10 Negri, J., Menon, V. & Young-Pearse, T. L. Assessment of Spontaneous Neuronal Activity In Vitro Using Multi-Well Multi-Electrode Arrays: Implications for Assay Development. eNeuro 7, doi:10.1523/ENEURO.0080-19.2019 (2020).

11 Romoli, M., Sen, A., Parnetti, L., Calabresi, P. & Costa, C. Amyloid-beta: a potential link between epilepsy and cognitive decline. Nat Rev Neurol 17, 469–485, doi:10.1038/s41582-021-00505-9 (2021).

12. Lomoio, S. et al. A 3D bioengineered neural tissue model generated from patient-derived iPSCs develops Alzheimer‘s disease-related phenotypes. 2022.2007.2021.501004, doi:10.1101/2022.07.21.501004 %J bioRxiv (2022).

13 Targa Dias Anastacio, H., Matosin, N. & Ooi, L. Neuronal hyperexcitability in Alzheimer’s disease: what are the drivers behind this aberrant phenotype? Transl Psychiatry 12, 257, doi:10.1038/s41398-022-02024-7 (2022).

14 Celone, K. A. et al. Alterations in memory networks in mild cognitive impairment and Alzheimer’s disease: an independent component analysis. J Neurosci 26, 10222–10231, doi:10.1523/jneurosci.2250-06.2006 (2006).

15 Dickerson, B. C. et al. Increased hippocampal activation in mild cognitive impairment compared to normal aging and AD. Neurology 65, 404–411, doi:10.1212/01.wnl.0000171450.97464.49 (2005).

16 Bookheimer, S. Y. et al. Patterns of brain activation in people at risk for Alzheimer’s disease. The New England journal of medicine 343, 450–456, doi:10.1056/nejm200008173430701 (2000).

17 Amatniek, J. C. et al. Incidence and predictors of seizures in patients with Alzheimer’s disease. Epilepsia 47, 867–872, doi:10.1111/j.1528-1167.2006.00554.x (2006).

18 Mendez, M. F., Catanzaro, P., Doss, R. C. R. A. R. & Frey, W. H., 2nd. Seizures in Alzheimer’s disease: clinicopathologic study. Journal of geriatric psychiatry and neurology 7, 230–233, doi:10.1177/089198879400700407 (1994).

19 Giorgio, J., Adams, J. N., Maass, A., Jagust, W. J. & Breakspear, M. Amyloid induced hyperexcitability in default mode network drives medial temporal hyperactivity and early tau accumulation. Neuron, 10.1016/j.neuron.2023.11.014 (2023).

20 Yang, F. et al. Alzheimer’s disease and epilepsy: An increasingly recognized comorbidity. Frontiers in aging neuroscience 14, 940515, doi:10.3389/fnagi.2022.940515 (2022).

21 Kazim, S. F. et al. Neuronal Network Excitability in Alzheimer’s Disease: The Puzzle of Similar versus Divergent Roles of Amyloid β and Tau. eNeuro 8, doi:10.1523/eneuro.0418-20.2020 (2021).

22 Bassett, S. S. et al. Familial risk for Alzheimer’s disease alters fMRI activation patterns. Brain 129, 1229–1239, doi:10.1093/brain/awl089 (2006).

23 McDonough, I. M. et al. Young Adults with a Parent with Dementia Show Early Abnormalities in Brain Activity and Brain Volume in the Hippocampus: A Matched Case-Control Study. Brain Sci 12, doi:10.3390/brainsci12040496 (2022).

24 Calafate, S. et al. Early alterations in the MCH system link aberrant neuronal activity and sleep disturbances in a mouse model of Alzheimer’s disease. Nature neuroscience, doi:10.1038/s41593-023-01325-4 (2023).

25 Busche, M. A. et al. Critical role of soluble amyloid-β for early hippocampal hyperactivity in a mouse model of Alzheimer’s disease. Proc Natl Acad Sci U S A 109, 8740–8745, doi:10.1073/pnas.1206171109 (2012).

26 Minkeviciene, R. et al. Amyloid beta-induced neuronal hyperexcitability triggers progressive epilepsy. J Neurosci 29, 3453–3462, doi:10.1523/jneurosci.5215-08.2009 (2009).

27 Bezprozvanny, I. & Mattson, M. P. Neuronal calcium mishandling and the pathogenesis of Alzheimer’s disease. Trends in neurosciences 31, 454–463, doi:10.1016/j.tins.2008.06.005 (2008).

28 Walton, H. S. & Dodd, P. R. Glutamate-glutamine cycling in Alzheimer’s disease. Neurochemistry international 50, 1052–1066, doi:10.1016/j.neuint.2006.10.007 (2007).

29 Gail Canter, R., et al. 3D mapping reveals network-specific amyloid progression and subcortical susceptibility in mice. Communications Biology 2, 360, doi:10.1038/s42003-019-0599-8 (2019).

30 Cunha, R. A. How does adenosine control neuronal dysfunction and neurodegeneration? J Neurochem 139, 1019–1055, doi:10.1111/jnc.13724 (2016).

31 Boison, D. Adenosine kinase: exploitation for therapeutic gain. Pharmacological reviews 65, 906–943, doi:10.1124/pr.112.006361 (2013).

32 Trinh, P. N. H., Baltos, J. A., Hellyer, S. D., May, L. T. & Gregory, K. J. Adenosine receptor signalling in Alzheimer’s disease. Purinergic Signal 18, 359–381, doi:10.1007/s11302-022-09883-1 (2022).

33 Cellai, L. et al. The Adenosinergic Signaling: A Complex but Promising Therapeutic Target for Alzheimer’s Disease. Front Neurosci 12, 520, doi:10.3389/fnins.2018.00520 (2018).

34 Lee, C. C. et al. Adenosine Augmentation Evoked by an ENT1 Inhibitor Improves Memory Impairment and Neuronal Plasticity in the APP/PS1 Mouse Model of Alzheimer’s Disease. Mol Neurobiol 55, 8936–8952, doi:10.1007/s12035-018-1030-z (2018).

35 Chang, C. P. et al. Equilibrative nucleoside transporter 1 inhibition rescues energy dysfunction and pathology in a model of tauopathy. Acta Neuropathol Commun 9, 112, doi:10.1186/s40478-021-01213-7 (2021).

36 Criscuolo, C. et al. Entorhinal Cortex dysfunction can be rescued by inhibition of microglial RAGE in an Alzheimer’s disease mouse model. Scientific Reports 7, 42370, doi:10.1038/srep42370 (2017).

37 Chistiakova, M., Bannon, N. M., Bazhenov, M. & Volgushev, M. Heterosynaptic plasticity: multiple mechanisms and multiple roles. The Neuroscientist : a review journal bringing neurobiology, neurology and psychiatry 20, 483–498, doi:10.1177/1073858414529829 (2014).

38 Pascual, O. et al. Astrocytic purinergic signaling coordinates synaptic networks. Science 310, 113–116, doi:10.1126/science.1116916 (2005).

39 Sun, Q. et al. Frequency-Dependent Synaptic Dynamics Differentially Tune CA1 and CA2 Pyramidal Neuron Responses to Cortical Input. 41, 8103–8110, doi:10.1523/JNEUROSCI.0451-20.2021%J The Journal of Neuroscience (2021).

40 Halassa, M. M. et al. Astrocytic modulation of sleep homeostasis and cognitive consequences of sleep loss. Neuron 61, 213–219, doi:10.1016/j.neuron.2008.11.024 (2009).

41 Williams-Karnesky, R. L. et al. Epigenetic changes induced by adenosine augmentation therapy prevent epileptogenesis. J Clin Invest 123, 3552–3563, doi:10.1172/JCI65636 (2013).

42 Murano, C. et al. Effect of the ketogenic diet in excitable tissues. Am J Physiol Cell Physiol 320, C547–C553, doi:10.1152/ajpcell.00458.2020 (2021).

43 Hertz, L., Chen, Y. & Waagepetersen, H. S. Effects of ketone bodies in Alzheimer’s disease in relation to neural hypometabolism, beta-amyloid toxicity, and astrocyte function. J Neurochem 134, 7–20, doi:10.1111/jnc.13107 (2015).

44 Elamin, M., Ruskin, D. N., Sacchetti, P. & Masino, S. A. A unifying mechanism of ketogenic diet action: The multiple roles of nicotinamide adenine dinucleotide. Epilepsy Res 167, 106469, doi:10.1016/j.eplepsyres.2020.106469 (2020).

45 Milstein, Aaron D. et al. Inhibitory Gating of Input Comparison in the CA1 Microcircuit. Neuron 87, 1274–1289, 10.1016/j.neuron.2015.08.025 (2015).

46 Sloin, H. E. et al. Local activation of CA1 pyramidal cells induces theta-phase precession. Science 383, 551–558, doi:10.1126/science.adk2456 (2024).

47 Jones, D. T. et al. Cascading network failure across the Alzheimer’s disease spectrum. Brain 139, 547–562, doi:10.1093/brain/awv338 (2016).

48 Raut, S. et al. Hypometabolism, Alzheimer’s Disease, and Possible Therapeutic Targets: An Overview. Cells 12, doi:10.3390/cells12162019 (2023).

49 Neuner, S. M., Wilmott, L. A., Hoffmann, B. R., Mozhui, K. & Kaczorowski, C. C. Hippocampal proteomics defines pathways associated with memory decline and resilience in normal aging and Alzheimer’s disease mouse models. Behavioural brain research 322, 288–298, doi:10.1016/j.bbr.2016.06.002 (2017).

50 Dennissen, F. J., Anglada-Huguet, M., Sydow, A., Mandelkow, E. & Mandelkow, E. M. Adenosine A1 receptor antagonist rolofylline alleviates axonopathy caused by human Tau ΔK280. Proc Natl Acad Sci U S A 113, 11597–11602, doi:10.1073/pnas.1603119113 (2016).

51 Zhou, L.-T. et al. Tau pathology epigenetically remodels the neuron-glial cross-talk in Alzheimer’s disease. 9, eabq7105, doi:doi:10.1126/sciadv.abq7105 (2023).

52 Orr, A. G. et al. Astrocytic adenosine receptor A2A and Gs-coupled signaling regulate memory. Nature neuroscience 18, 423–434, doi:10.1038/nn.3930 (2015).

53 Goncalves, F. Q. et al. Synaptic and memory dysfunction in a beta-amyloid model of early Alzheimer’s disease depends on increased formation of ATP-derived extracellular adenosine. Neurobiol Dis 132, 104570, doi:10.1016/j.nbd.2019.104570 (2019).

54 Lopes, C. R. et al. Adenosine A2A Receptor Up-Regulation Pre-Dates Deficits of Synaptic Plasticity and of Memory in Mice Exposed to Aβ1–42 to Model Early Alzheimer’s Disease. 13, 1173 (2023).

55 Lee, K. S., Schubert, P., Reddington, M. & Kreutzberg, G. W. The distribution of adenosine A1 receptors and 5’-nucleotidase in the hippocampal formation of several mammalian species. The Journal of comparative neurology 246, 427–434, doi:10.1002/cne.902460402 (1986).

56 Muñoz, M.-D. & Solís, J. M. Characterisation of the mechanisms underlying the special sensitivity of the CA2 hippocampal area to adenosine receptor antagonists. Neuropharmacology 144, 9–18, 10.1016/j.neuropharm.2018.10.017 (2019).

57 Badimon, A. et al. Negative feedback control of neuronal activity by microglia. Nature 586, 417–423, doi:10.1038/s41586-020-2777-8 (2020).

58 Wang, Y. et al. TREM2 lipid sensing sustains the microglial response in an Alzheimer’s disease model. Cell 160, 1061–1071, doi:10.1016/j.cell.2015.01.049 (2015).

59 Krasemann, S. et al. The TREM2-APOE Pathway Drives the Transcriptional Phenotype of Dysfunctional Microglia in Neurodegenerative Diseases. Immunity 47, 566–581.e569, doi:10.1016/j.immuni.2017.08.008 (2017).

60 Murugan, M. et al. Adenosine kinase: An epigenetic modulator in development and disease. Neurochemistry international 147, 105054, 10.1016/j.neuint.2021.105054 (2021).

61 Boison, D. Adenosine dysfunction and adenosine kinase in epileptogenesis. The open neuroscience journal 4, 93–101, doi:10.2174/1874082001004020093 (2010).

62 Wu, Y. et al. BHBA treatment improves cognitive function by targeting pleiotropic mechanisms in transgenic mouse model of Alzheimer’s disease. FASEB J 34, 1412–1429, doi:10.1096/fj.201901984R (2020).

63 Shippy, D. C., Wilhelm, C., Viharkumar, P. A., Raife, T. J. & Ulland, T. K. beta-Hydroxybutyrate inhibits inflammasome activation to attenuate Alzheimer’s disease pathology. J Neuroinflammation 17, 280, doi:10.1186/s12974-020-01948-5 (2020).

64 Pawlosky, R. J., Kashiwaya, Y., King, M. T. & Veech, R. L. A Dietary Ketone Ester Normalizes Abnormal Behavior in a Mouse Model of Alzheimer’s Disease. Int J Mol Sci 21, doi:10.3390/ijms21031044 (2020).

65 Krishnan, M. et al. beta-hydroxybutyrate Impedes the Progression of Alzheimer’s Disease and Atherosclerosis in ApoE-Deficient Mice. Nutrients 12, doi:10.3390/nu12020471 (2020).

66 Saito, T. et al. Single App knock-in mouse models of Alzheimer’s disease. Nature neuroscience 17, 661–663, doi:10.1038/nn.3697 (2014).

67 Xiong, G., Metheny, H., Johnson, B. N. & Cohen, A. S. A Comparison of Different Slicing Planes in Preservation of Major Hippocampal Pathway Fibers in the Mouse. 11, doi:10.3389/fnana.2017.00107 (2017).

68 Sadick, J. S. et al. Astrocytes and oligodendrocytes undergo subtype-specific transcriptional changes in Alzheimer’s disease. Neuron 110, 1788–1805 e1710, doi:10.1016/j.neuron.2022.03.008 (2022).

