## Supplementary Material and Figures for "Adenosine deficiency facilitates CA1 synaptic hyperexcitability in the presymptomatic phase of a knock in mouse model of Alzheimer’s disease"

[#] equally contributed

**Table S1. Statistical Analysis.** Statistical details summarized by figure panels. In bold, statistically significant variables.

| Figure | Dependent Variable | Independent Variable | Statistic | |
|---|---|---|---|---|
| 1B | fEPSP-FV slope | Genotype: WT, APPKI<br>Sex: Male, Female | **Genotype**<br>Sex<br>Genotype*Sex | **F(1,30.5)=4.58, p=0.04**<br>F(1,30.5)=7E-4, p=0.98<br>F(1,30.5)=0.02, p=0.90 |
| 1D | fEPSP-FV slope | Genotype: WT, APPKI<br>Sex: Male, Female<br>Drug: Pre, Post | Genotype<br>**Drug**<br>Sex<br>**Genotype*Drug**<br>Genotype*Sex<br>Drug*Sex<br>Genotype*Drug*Sex | F(1,3.90)=0.33, p=0.60<br>**F(1,12)=21.66, p=5.5E-4**<br>F(1,3.90)=0.76, p=0.43<br>**F(1,12)=8.51, p=0.013**<br>F(1,3.90)=3.19, p=0.15<br>F(1,12)=0.77, p=0.40<br>F(1,12)=3.63, p=0.08 |
| 1F | fEPSP-FV slope | Genotype: WT, APPKI<br>Sex: Male, Female | **Genotype**<br>Sex<br>Genotype*Sex | **F(1,17.2)=6.38, p=0.022**<br>F(1,17.2)=4E-4, p=0.98<br>F(1,17.2)=1.13, p=0.30 |
| 1H | fEPSP-FV slope | Genotype: WT, APPKI<br>Sex: Male, Female<br>Drug: Pre, Post | **Genotype**<br>Drug<br>Sex<br>Genotype*Drug<br>Genotype*Sex<br>Drug*Sex<br>Genotype*Drug*Sex | **F(1,9)=5.30, p=0.05**<br>F(1,9)=1.47, p=0.26<br>F(1,9)=0.15, p=0.71<br>F(1,9)=0.14, p=0.72<br>F(1,9)=0.58, p=0.47<br>F(1,9)=0.27, p=0.61<br>F(1,9)=0.07, p=0.79 |
| 2C | Norm. fEPSP | Genotype: WT, APPKI<br>Time: Pre, Post<br>Sex: Male, Female | Genotype<br>**Time**<br>Sex<br>Genotype*Time<br>Genotype*Sex<br>Time*Sex<br>Genotype*Time*Sex | F(1,8)=0.11, p=0.92<br>**F(1,8)=92.25, p=1.1e-5**<br>F(1,8)=0.33, p=0.58<br>F(1,8)=0.01, p=0.92<br>F(1,8)=3.02, p=0.13<br>F(1,8)=0.33, p=0.58<br>F(1,8)=3.02, p=0.12 |
| 2I | Norm. fEPSP S1 | Genotype: WT, APPKI<br>Sex: Male, Female<br>Time: S1, S2-S1 | Genotype<br>Time<br>Sex<br>Genotype*Time<br>Genotype*Sex<br>Time*Sex<br>Genotype*Time*Sex | F(1,16)=1.32, p=0.27<br>F(1,16)=1.05, p=0.32<br>F(1,16)=0.09, p=0.77<br>F(1,16)=1.32, p=0.27<br>F(1,16)=0.63, p=0.44<br>F(1,16)=0.09, p=0.77<br>F(1,16)=0.63, p=0.44 |
| 2J | Norm. fEPSP S2 | Genotype: WT, APPKI<br>Sex: Male, Female<br>Time: Pre, Post | Genotype<br>**Time**<br>Sex<br>Genotype*Time<br>Genotype*Sex<br>Time*Sex<br>Genotype*Time*Sex | F(1,7)=1.06, p=0.34<br>**F(1,7)=179.46, p=3E-6**<br>F(1,7)=0.33, p=0.58<br>F(1,7)=3.28, p=0.11<br>F(1,7)=3.39, p=0.11<br>F(1,7)=1.22, p=0.31<br>F(1,7)=0.58, p=0.47 |
| 2K | Norm. fEPSP S1 | Genotype: WT, APPKI<br>Sex: Male, Female<br>Time: Pre, Post | **Genotype**<br>**Time**<br>Sex<br>**Genotype*Time**<br>Genotype*Sex<br>Time*Sex<br>**Genotype*Time*Sex** | **F(1,7.5)=11.52, p=0.01**<br>**F(1,7.1)=93.28, p=2e-5**<br>F(1,7.5)=0.50, p=0.50<br>**F(1,7.1)=27.46, p=1e-3**<br>F(1,7.5)=1.41, p=0.27<br>F(1,7.1)=0.02, p=0.88<br>**F(1,7.1)=11.701, p=0.01** |
| 3C | PS Amplitude-FV slope | Genotype: WT, APPKI<br>Sex: Male, Female | **Genotype**<br>Sex<br>Genotype*Sex | **F(1,24)=4.24, p=0.05**<br>F(1,24)=0.02, p=0.88<br>F(1,24)=0.68, p=0.42 |
| 3E | PS Amplitude-FV slope | Genotype: WT, APPKI<br>Sex: Male, Female<br>Drug: Pre, Post | Genotype<br>**Drug**<br>Sex<br>Genotype*Drug<br>Genotype*Sex<br>Drug*Sex<br>**Genotype*Drug*Sex** | F(1,6.82)=0.79, p=0.41<br>**F(1,10.4)=41.25, p=6E-5**<br>F(1,7.1)=0.10, p=0.76<br>F(1,10.4)=2.19, p=0.17<br>F(1,7.1)=0.42, p=0.54<br>F(1,11.5)=0.06, p=0.81<br>**F(1,11.5)=6.23, p=0.03** |
| 3F | PS Amplitude-fEPSP slope | Genotype: WT, APPKI<br>Sex: Male, Female | Genotype<br>Sex<br>Genotype*Sex | F(1,24)=0.08, p=0.78<br>F(1,24)=0.32, p=0.58<br>F(1,24)=0.54, p=0.47 |
| 3H | PS Amplitude-fEPSP slope | Genotype: WT, APPKI<br>Sex: Male, Female<br>Drug: Pre, Post | Genotype<br>**Drug**<br>Sex | F(1,10.80)=0.16, p=0.70<br>**F(1,10.39)=8.19, p=0.02**<br>F(1,15.41)=0.14, p=0.71 |

| | | | | |
|---|---|---|---|---|
| | | | Genotype*Drug<br>Genotype*Sex<br>Drug*Sex<br>Genotype*Drug*Sex | F(1,10.39)=0.81, p=0.39<br>F(1,15.41)=0.11, p=0.75<br>F(1,10.23)=0.52, p=0.49<br>F(1,10.23)=0.41, p=0.54 |
| 3L | *P*(PS) | Condition: WT, WT+CPT, APPKI, APPKI+CPT<br>Sex: Male, Female | Condition<br>Sex<br>Condition*Sex | F(3,18)=1.64, p=0.22<br>F(1,18)=3E-4, p=0.99<br>F(3,18)=1.70, p=0.20 |
| 3L | *P*(PS) | Condition: WT, WT+CPT, APPKI, APPKI+CPT<br>Band: High, Low | Condition<br>**Band**<br>**Condition*Band** | F(3,24.19)=3.01, p=0.06<br>**F(1,9.4)=21.76, p=0.001**<br>F**(3,9.4)=4.08, p=0.04** |
| 4B | Adenosine dialysate | Genotype: WT, APPKI<br>Sex: Male, Female | **Condition**<br>Sex<br>Condition*Sex | **F(1,8)=7.62, p=0.03**<br>F(1,8)=0.01, p=0.93<br>F(1,8)=2.48, p=0.15 |
| 4F | CD39 | Genotype: WT, APPKI<br>Sex: Male, Female | **Genotype**<br>Sex<br>Genotype*Sex | **F(1,11)=8.59, p=0.014**<br>F(1,11)=2e-4, p=0.99<br>F(1,11)=0.59, p=0.46 |
| | CD73 | Genotype: WT, APPKI<br>Sex: Male, Female | Genotype<br>Sex<br>Genotype*Sex | F(1,14)=4.39, p=0.055<br>F(1,14)=0.005 p=0.94<br>F(1,14)=0.02, p=0.88 |
| | ENT1 | Genotype: WT, APPKI<br>Sex: Male, Female | Genotype<br>Sex<br>Genotype*Sex | F(1,11)=0.61, p=0.45<br>F(1,11)=1.92, p=0.19<br>F(1,11)=1.59, p=0.23 |
| | SAHH | Genotype: WT, APPKI<br>Sex: Male, Female | Genotype<br>Sex<br>Genotype*Sex | F(1,15)=3E-3, p=0.95<br>F(1,15)=2E-2 p=0.96<br>F(1,15)=0.02 p=0.88 |
| | ADA | Genotype: WT, APPKI<br>Sex: Male, Female | Genotype<br>Sex<br>Genotype*Sex | F(1,17)=2.83, p=0.11<br>F(1,17)=0.71, p=0.41<br>F(1,17)=0.004, p=0.95 |
| | ADK-L | Genotype: WT, APPKI<br>Sex: Male, Female | **Genotype**<br>Sex<br>Genotype*Sex | **F(1,14)=17.03, p=1E-3**<br>F(1,14)=0.01, p=0.94<br>F(1,14)=0.04, p=0.83 |
| | ADK-S | Genotype: WT, APPKI<br>Sex: Male, Female | Genotype<br>Sex<br>Genotype*Sex | F(1,14)=0.02, p=0.88<br>F(1,14)=0.43, p=0.52<br>F(1,14)=0.05, p=0.82 |
| | $A_1R$ | Genotype: WT, APPKI<br>Sex: Male, Female | Genotype<br>Sex<br>Genotype*Sex | F(1,8)=0.03, p=0.87<br>F(1,8)=0.77, p=0.40<br>F(1,8)=3.30, p=0.11 |
| 4I | fEPSP-FV slope | Genotype: WT, APPKI<br>Sex: Male, Female<br>Drug: Pre, Post | Genotype<br>**Drug**<br>Sex<br>Genotype*Drug<br>Genotype*Sex<br>Drug*Sex<br>Genotype*Drug*Sex | F(1,10)=2.85, p=0.12<br>**F(1,10)=5.16, p=0.04**<br>F(1,10)=0.01, p=0.90<br>F(1,10)=4.12, p=0.07<br>F(1,10)=0.06, p=0.81<br>F(1,10)=0.16, p=0.70<br>F(1,10)=6e-3, p=0.94 |
| 5B-Left | fEPSP-FV slope | Treatment: DMSO, 5-ITU<br>Sex: Male, Female | **Treatment**<br>Sex<br>Drug*Sex | **F(1,12)=26.98, p=2e-4**<br>F(1,12)=0.06, p=0.81<br>F(1,12)=1.19, p=0.30 |
| 5B-Right | fEPSP-FV slope | Treatment: DMSO, 5-ITU<br>Sex: Male, Female<br>Drug: Pre, Post | Treatment<br>Drug<br>Sex<br>Treatment*Drug<br>Genotype*Sex<br>Drug*Sex<br>Treatment *Drug*Sex | **F(1,11.7)=59.21, p=6e-6**<br>**F(1,9.01)=7.96, p=0.02**<br>F(1,11.7)=4e-3, p=0.94<br>**F(1,9.01)=8.61, p=0.02**<br>F(1,11.7)=3.67, p=0.08<br>F(1,9.01)=4e-3, p=0.94<br>F(1,9.01)=0.08, p=0.78 |
| 5D-Left | FV-Pop. Spike slope | Treatment: DMSO, 5-ITU<br>Sex: Male, Female | **Treatment**<br>Sex<br>Drug*Sex | **F(1,6)=6.56, p=0.04**<br>F(1,6)=2.92, p=0.14<br>F(1,6)=1.34, p=0.29 |
| 5D-right | FV-Pop. Spike slope | Treatment: DMSO, 5-ITU<br>Sex: Male, Female<br>Drug: Pre, Post | **Treatment**<br>**Drug**<br>Sex<br>Treatment*Drug<br>Genotype*Sex<br>Drug*Sex<br>Treatment *Drug*Sex | **F(1,7)=6.85, p=0.03**<br>**F(1,7)=34.31, p=6e-4**<br>F(1,7)=0.04, p=0.84<br>F(1,7)=2.24, p=0.18<br>F(1,7)=1.99, p=0.20<br>F(1,7)=0.06, p=0.81<br>F(1,7)=3.63, p=0.10 |
| 5E | CD39 | Treatment: DMSO, 5-ITU<br>Sex: Male, Female | Treatment<br>Sex<br>Drug*Sex | F(1,7)=0.15, p=0.71<br>F(1,7)=1.05, p=0.34<br>F(1,7)=0.26, p=0.63 |
| | | Treatment: DMSO, 5-ITU | **Treatment** | **F(1,6)=6.86, p=0.04** |

| | | | | |
|---|---|---|---|---|
| | CD73 | Sex: Male, Female | Sex<br>Drug*Sex | F(1,6)=1.56, p=0.26<br>F(1,6)=1e-3, p=0.92 |
| 5G | Weight | Diet: RD, KD<br>Time: d0/d7/d14/d21/d28<br>Sex: Male, Female | **Diet**<br>**Day**<br>**Sex**<br>**Diet*Day**<br>**Diet*Sex**<br>**Day*Sex**<br>Diet*Day*Sex | **F(1,34.5)=155.2, p=3e-14**<br>**F(4,31.9)=8.90, p=6e-5**<br>**F(1,34.5)=16.54, p=3e-4**<br>**F(4,31.9)=56.83, p=4e-14**<br>**F(1,34.5)=11.79, p=2e-3**<br>**F(4,31.9)=4.42, p=6e-3**<br>F(4,31.9)=2.13, p=0.10 |
| 5H | Ketone Bodies/Weight | Diet: RD, KD<br>Time: d0/d7/d14/d21/d28<br>Sex: Male, Female | **Diet**<br>**Day**<br>Sex<br>**Diet*Day**<br>**Diet*Sex**<br>Day*Sex<br>Diet*Day*Sex | **F(1,14)=76.64, p=5e-7**<br>**F(4,14)=30.19, p=9e-7**<br>F(1,14)=4.49, p=0.052<br>**F(4,14)=33.68, p=5e-7**<br>**F(1,14)=4.64, p=0.05**<br>F(4,14)=2.49, p=0.1<br>F(4,14)=2.31, p=0.11 |
| 5I | Glucose/Weight | Diet: RD, KD<br>Time: d0/d7/d14/d21/d28<br>Sex: Male, Female | **Diet**<br>**Day**<br>**Sex**<br>**Diet*Day**<br>Diet*Sex<br>Day*Sex<br>**Diet*Day*Sex** | **F(1,12.6)=23.29, p=4e-4**<br>**F(4,11.4)=58.42, p=2e-7**<br>F(1,12.6)=0.09, p=0.76<br>**F(4,11.4)=16.84, p=1e-4**<br>F(1,12.6)=2.17, p=0.16<br>F(4,11.4)=0.64, p=0.64<br>**F(4,11.4)=6.69, p=5e-3** |
| 5K-Left | fEPSP-FV slope | Diet: RD, KD<br>Sex: Male, Female | **Diet**<br>Sex<br>Diet*Sex | **F(1,15.9)=10.05, p=0.006**<br>F(1,15.9)=1.46, p=0.25<br>F(1,15.9)=1.57, p=0.23 |
| 5K-Right | fEPSP-FV slope | Diet: RD, KD<br>Sex: Male, Female<br>Drug: Pre, Post | Genotype<br>**Drug**<br>Sex<br>**Genotype*Drug**<br>Genotype*Sex<br>Drug*Sex<br>Genotype*Drug*Sex | F(1,14)=2.52, p=0.14<br>**F(1,14)=20.27, p=5e-4**<br>F(1,14)=0.56, p=0.46<br>**F(1,14)=14.08, p=0.002**<br>F(1,14)=0.40, p=0.54<br>F(1,14)=2.02, p=0.18<br>F(1,14)=0.003, p=0.96 |
| 5M-Left | FV-Pop. Spike slope | Diet: RD, KD<br>Sex: Male, Female | **Diet**<br>Sex<br>Diet*Sex | **F(1,9)=5.18, p=0.05**<br>F(1,9)=0.05, p=0.82<br>F(1,9)=0.37, p=0.56 |
| 5M-Right | FV-PS slope | Diet: RD, KD<br>Sex: Male, Female<br>Drug: Pre, Post | Genotype<br>**Drug**<br>Sex<br>Genotype*Drug<br>Genotype*Sex<br>Drug*Sex<br>Genotype*Drug*Sex | F(1,7)=2.54, p=0.15<br>**F(1,7)=7.86, p=0.03**<br>F(1,7)=0.43, p=0.53<br>F(1,7)=2.41, p=0.16<br>F(1,7)=0.48, p=0.51<br>F(1,7)=0.79, p=0.40<br>F(1,7)=0.76, p=0.41 |
| 5N | CD39 | Diet: RD, KD<br>Sex: Male, Female | Diet<br>Sex<br>**Diet*Sex** | F(1,18)=2.36, p=0.14<br>F(1,18)=2.49, p=0.13<br>**F(1,18)=4.55, p=0.05** |
| | CD73 | Diet: RD, KD<br>Sex: Male, Female | **Diet**<br>Sex<br>Diet*Sex | **F(1,18)=12.52, p=2e-3**<br>F(1,18)=3.94, p=0.06<br>F(1,18)=1.28, p=0.27 |
| S1B | Stimulation-FV slope | Genotype: WT, APPKI<br>Sex: Male, Female | Genotype<br>Sex<br>Genotype*Sex | F(1,28.7)=0.13, p=0.72<br>F(1,28.7)=0.64, p=0.21<br>F(1,28.7)=0.13, p=0.73 |
| S1D | Stimulation -FV slope | Genotype: WT, APPKI<br>Sex: Male, Female | Genotype<br>Sex<br>Genotype*Sex | F(1,30)=0.36, p=0.55<br>F(1,30)=0.49, p=0.49<br>F(1,30)=0.01, p=0.94 |
| S2D | fEPSP-FV slope | Genotype: WT, APPKI<br>Sex: Male, Female<br>Drug: Pre, Post | **Genotype**<br>**Drug**<br>Sex<br>Genotype*Drug<br>Genotype*Sex<br>Drug*Sex<br>Genotype*Drug*Sex | **F(1,6)=9.07, p=0.03**<br>**F(1,6)=33.87, p=1E-3**<br>F(1,6)=1.50, p=0.27<br>F(1,6)=4.97, p=0.07<br>F(1,6)=0.40, p=0.55<br>F(1,6)=0.63, p=0.46<br>F(1,6)=0.28, p=0.62 |
| S2H | fEPSP-FV slope | Genotype: WT, APPKI<br>Sex: Male, Female<br>Drug: Pre, Post | **Genotype**<br>**Drug**<br>**Sex**<br>Genotype*Drug<br>**Genotype*Sex**<br>Drug*Sex | **F(1,6)=9.52, p=0.02**<br>**F(1,6)=29.86, p=0.002**<br>**F(1,6)=17.00, p=0.006**<br>F(1,6)=1.47, p=0.13<br>**F(1,6)=10.27, p=0.02**<br>F(1,6)=0.40, p=0.55 |

| | | | | |
|---|---|---|---|---|
| | | | Genotype*Drug*Sex | F(1,6)=0.43, p=0.53 |
| S3C | fEPSP-FV slope | Drug: Baseline, 5-ITU, 5-IT+CPT | **Drug** | **F(2)=17.41, p=0.02** |
| S4G | Phase 1st Pop. Spike | Condition: WT, WT+CPT, APPKI, APPKI+CPT<br>Sex: Male, Female | Condition<br>Sex<br>Condition*Sex | F(3,28)=0.98, p=0.42<br>F(1,28)=0.18, p=0.68<br>F(3,28)=0.51, p=0.68 |
| S4H | Phase Adaptation Index | Condition: WT, WT+CPT, APPKI, APPKI+CPT<br>Sex: Male, Female | Condition<br>Sex<br>Condition*Sex | F(3,25)=0.20, p=0.89<br>F(1,25)=0.01, p=0.93<br>F(3,25)=0.47, p=0.71 |
| S4I | Frequency Adaptation Index | Condition: WT, WT+CPT, APPKI, APPKI+CPT<br>Sex: Male, Female | **Condition**<br>**Sex**<br>**Condition*Sex** | **F(3,26)=4.11, p=0.02**<br>**F(1,26)=5.39, p=0.03**<br>**F(3,26)=3.61, p=0.03** |
| S7C | fEPSP-FV slope | Genotype: hAPP, APPKI<br>Sex: Male, Female<br>Drug: Pre, Post | **Genotype**<br>**Drug**<br>Sex<br>**Genotype*Drug**<br>Genotype*Sex<br>Drug*Sex<br>Genotype*Drug*Sex | **F(1,7.3)=21.82, p=2e-3**<br>**F(1,14.9)=84.13, p=2e-7**<br>F(1,7.3)=2.33, p=0.17<br>**F(1,14.9)=53.64, p=3e-6**<br>F(1,7.3)=0.18, p=0.68<br>F(1,14.9)=1.93, p=0.18<br>F(1,14.9)=0.06, p=0.81 |
| S7E | PS-FV slope | Genotype: hAPP, APPKI<br>Sex: Male, Female | **Genotype**<br>Sex<br>Genotype*Sex | **F(1,16)=4.50, p=0.05**<br>F(1,16)=0.06, p=0.81<br>F(1,16)=0.28, p=0.60 |
| S7F | PS-FV slope | Genotype: hAPP, APPKI<br>Sex: Male, Female<br>Drug: Pre, Post | Genotype<br>**Drug**<br>Sex<br>Genotype*Drug<br>Genotype*Sex<br>Drug*Sex<br>Genotype*Drug*Sex | F(1,5.9)=0.23, p=0.65<br>**F(1,9)=23.49, p=9e-4**<br>F(1,5.9)=1e-3, p=0.96<br>F(1,9)=0.28, p=0.61<br>F(1,5.9)=0.09, p=0.77<br>F(1,9)=1.15, p=0.31<br>F(1,9)=0.98, p=0.34 |

**Supplementary Figures**

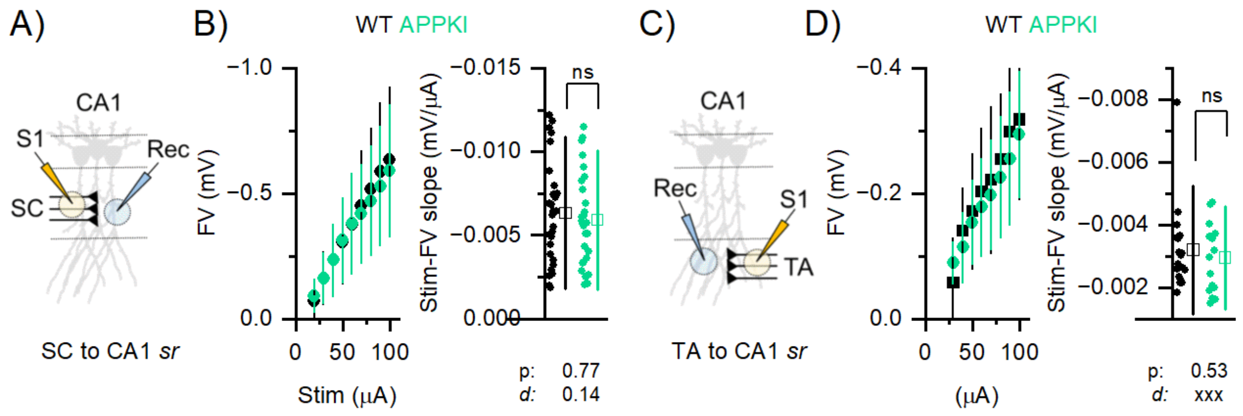


**Figure S1. Presynaptic fibers are not degenerated in the APPKI model.** We tested the hypothesis that the synaptic hyperactivity was caused by fiber degeneration of the SC and TA tracts. **A,C)** Scheme of experiment to monitor SC-to-CA1 and TA-to CA1 synaptic signaling (S1: stimulation; Rec: recording). **B,D)** Mean Stimulation Intensity-FV relationship (left) and individual slope data points (right); SC-to-CA1; WT: n=30/16, APPKI: n= 27/17; TA-to-CA1 - WT: n=18/11, APPKI: n= 16/9. The data indicated that, given the same stimulation intensity, the same fraction of presynaptic fibers (both SC and TA) have been activated in both genotypes supporting the absence of degeneration. p: p-value; d: Cohen's effect size. Mean±S.D. n=n. slices/n. mice. Statistic: generalized linear mixed effect models . Black: WT, Green: APPKI.

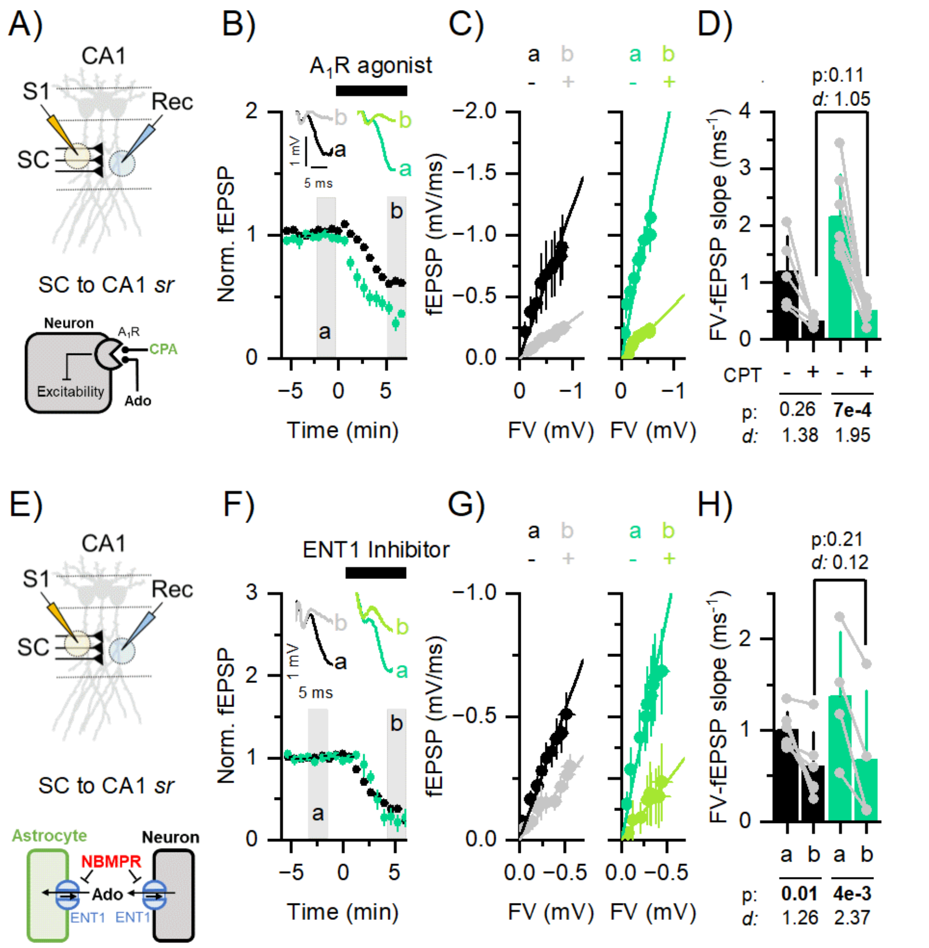


**Figure S2. A$_1$R-mediated and ENT1-mediated activities are intact in the APPKI model.** We tested the hypothesis that the synaptic hyperexcitability in the SC tract was mediated by either impaired A$_1$R activation and/or ENT1 functionality. **A,E)** Top: Scheme of experiment to monitor SC-to-CA1 synaptic signaling (S1: stimulation; Rec: recording). Bottom: cartoon of adenosine signaling. **B,F)** Time trajectory of normalized fEPSP before (a) and after (b) the dosage of the A$_1$R agonist, CPA (10 nM), or ENT1 inhibitor, NBMPR (300 nM). C,G) Mean fEPSP-F.V. relationship (left) and individual paired slope data points (right) before (a) and after (b) dosage of the A$_1$R agonist and ENT1 inhibitor; A1R agonist; WT: n=5/5, APPKI: n=7/7; ENT1 inhibitor - WT: n=6/6, APPKI: n=4/4. The data indicated that both A$_1$R and ENT1 signaling were intact in the APPKI model. p: p-value; d: Cohen's effect size. Mean±S.D. n=n. slices/n. mice. Statistic: generalized linear mixed effect models . Black: WT, Green: APPKI, Grey: WT+CPT, Light Green: APPKI+CPT.

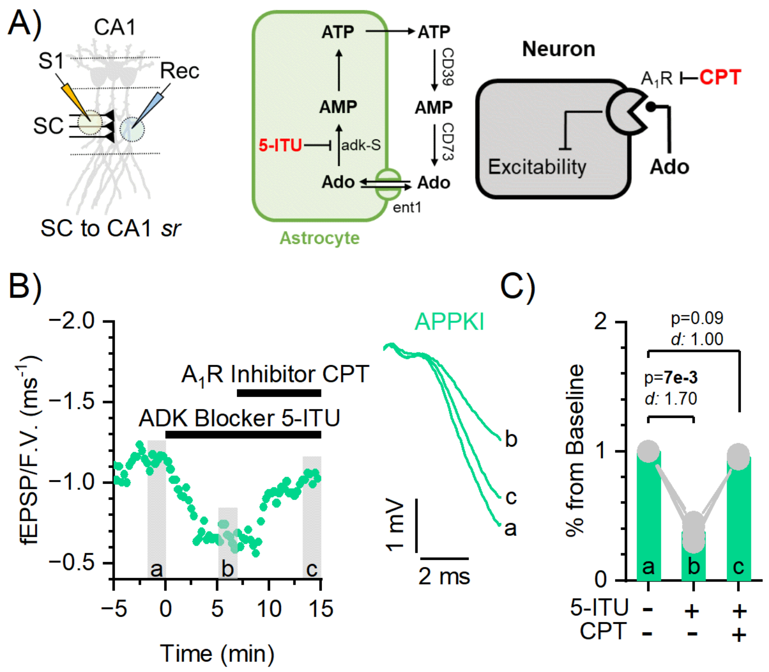


**Figure S3. Acute 5-ITU dosage in vitro increases extracellular concentrations of adenosine.** We tested the hypothesis that acute dosage of 5-ITU was able to increase extracellular adenosine by reducing the activity of ADK-S. Blocking ADK-S results in an elevation of intracellular adenosine levels, consequently causing a reversal in the flux direction of adenosine through the ENT1 transporter. This reversal leads to the efflux of adenosine from astrocytes rather than its influx. **A)** Left: Scheme of experiment to monitor SC-to-CA1 synaptic signaling (S1: stimulation; Rec: recording). Right: cartoon of adenosine signaling. **B)** Left: Time trajectory of normalized fEPSP/FV before (a), with 5-ITU (b) and 5-ITU+CPT (c). Right: Representative traces without drugs (a), with 5-ITU (b) and with 5-ITU+CPT. **C)** Individual paired slope data points at the three time points (a-c) as indicated; APPKI: n=4/4. The data indicated that the acute dosage of 5-ITU reduced synaptic activity in an $A_1R$-dependent way. It shows that the block of ADK is sufficient to elevate extracellular adenosine which, by actin through $A_1R$, depresses neuronal activity. Mean±S.D. n=n. slices/n. mice. Green: APPKI

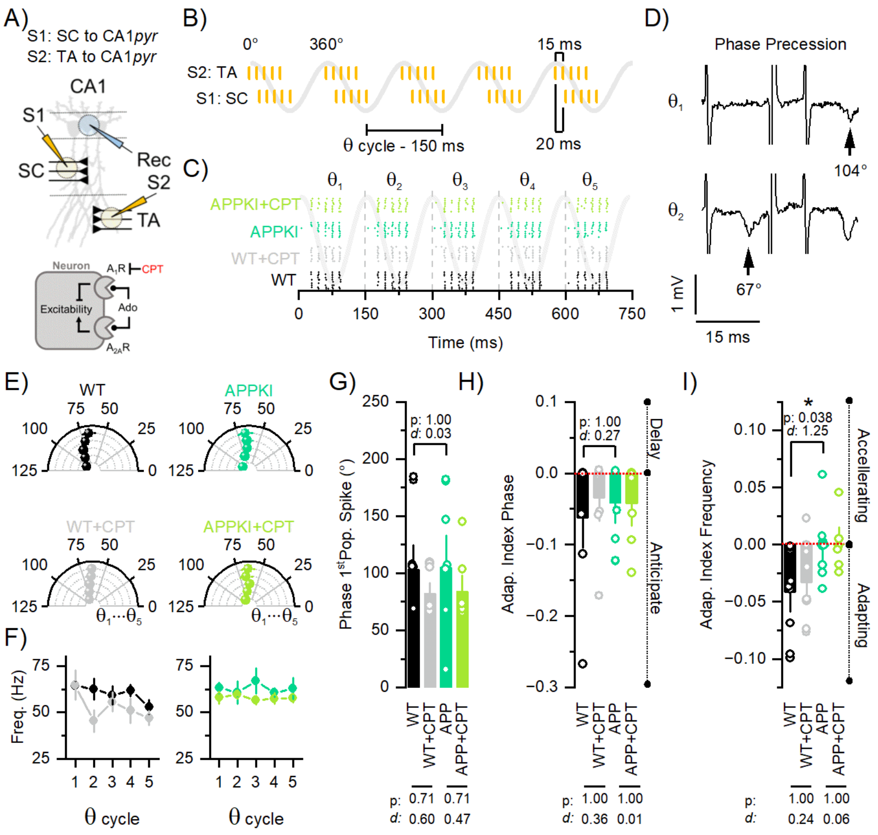


**Figure S4. Hippocampal circuit computation is minimally affected in the APPKI model.** CA1 output is shaped by the temporal patterns of stimulations occurring at the TA and SC terminals. With this temporal significance in mind, we aimed to investigate whether the altered synaptic transmission and adenosine deficiency beget circuit-level consequences. **A)** Top: Scheme of experiment to monitor SC-to-CA1 and TA-to CA1 synaptic signaling (S1: stimulation SC; S2: stimulation TA; Rec: recording). Bottom: cartoon of adenosine signaling. **B)** Theta stimulation protocol. Each theta cycle ( θ 1-5) is composed of 5 stimulations from each electrode (S2 and S1). The first S1 stimulation occurs 20 ms after the first stimulation from S2; within each electrode, the inter-stimulation interval is 15 ms (67 Hz). Each cycle is repeated every 150 ms (360°) five times. **C)** Raster plot of the time of population spike within each cycle and condition, as indicated. **D)** Representative traces showing the phase precession process, namely after the first cycle ( θ 1), the first population spike (arrow) is anticipated (from 104° to 67°) in phase. **E)** Mean phases of the first population spike at consecutive theta cycles for the different conditions, as indicated. **F)** Mean firing rate at consecutive theta cycles for the different conditions. **G)** Mean phases of the first population spike and individual data points (WT: n=10/10; WT+CPT: n=8/8; APPKI: n= 9/9; APPKI+CPT: n= 8/8). Mean adaptation indices of **H)** phase (WT: n=9/9; WT+CPT: n=8/8; APPKI: n= 9/9; APPKI+CPT: n= 7/7) and **I)** frequency (WT: n=10/10; WT+CPT: n=8/8; APPKI: n= 9/9; APPKI+CPT: n= 6/6) for the different conditions. These results demonstrated that when subjected to a naturalistic heterosynaptic stimulation engaging TA and SC terminals, the APPKI model exhibits an adenosine independent loss of frequency adaptation properties across theta cycles while preserving phase precession. p: p-value; d: Cohen's effect size. Mean±S.D. n=n. slices/n. mice. Statistic: generalized linear mixed effect models. Black: WT, Green: APPKI, Grey: WT+CPT, Light Green: APPKI+CPT.

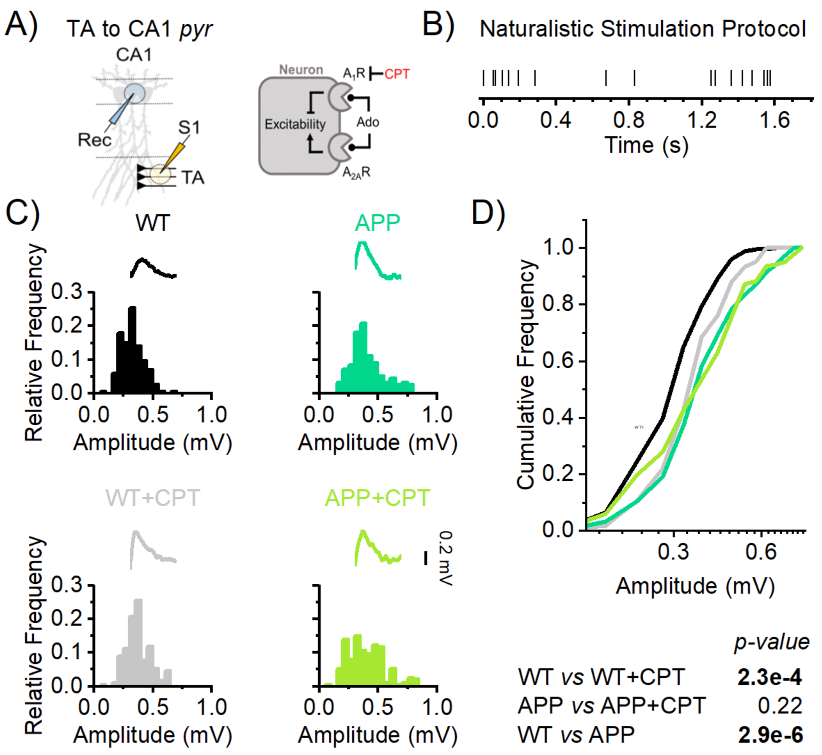


**Figure S5. Adenosine-dependent reduction of filtering properties in the APPKI model.** We tested the hypothesis that the synaptic signal generated at the TA terminals and recorded in the somatic layer exhibits reduced filtering in the APPKI model. **A)** Left: Scheme of experiment to monitor TA-to-CA1 somatic signaling (S1: stimulation; Rec: recording). Right: cartoon of adenosine signaling. **B)** Naturalistic stimulation protocol. **C)** Signal amplitude distributions in response to naturalistic stimulation protocol across conditions, as indicated (WT: n=6/4; WT+CPT: n=6/4; APPKI: n= 6/4; APPKI+CPT: n= 7/5). **D)** Top: Cumulative distribution reconstructed from the distributions. Bottom: Tabular results using Kolmogorov-Smirnov test. The data indicated that in the APPKI model the signal from the TA terminals to the CA1 soma underwent reduced filtering, resulting in larger somatic values. This effect was driven by reduced adenosine tone given that in the WT but not in the APPKI model the antagonism of the $A_1R$ reduced the filtering properties. Mean±S.D. n=n. slices/n. mice. Black: WT, Green: APPKI, Grey: WT+CPT, Light Green: APPKI+CPT.

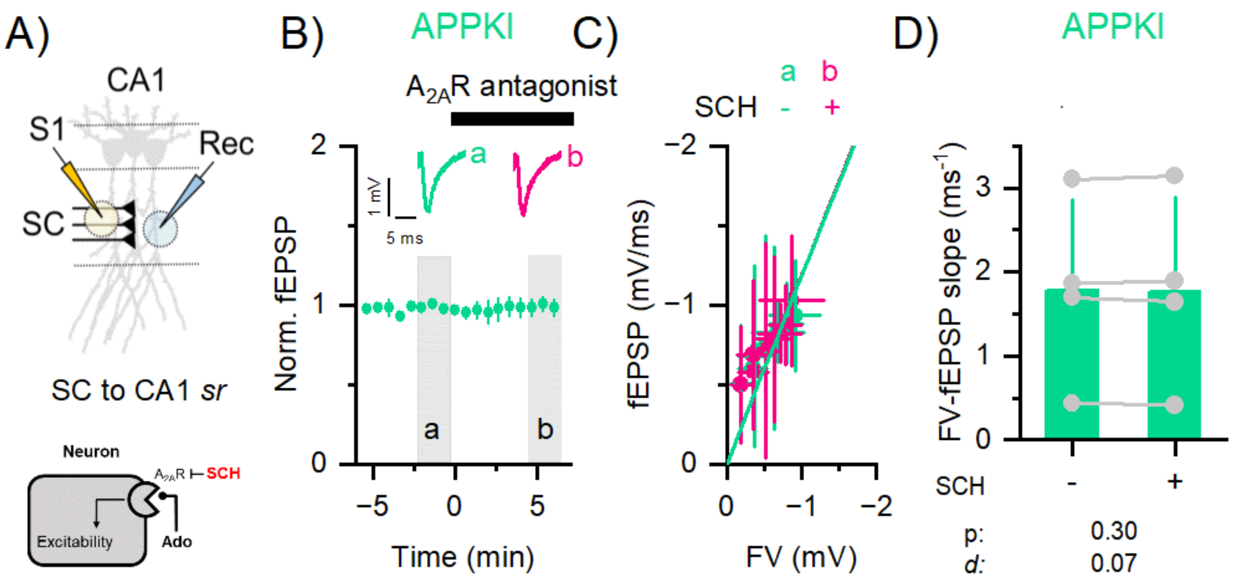


**Figure S6. Basal synaptic transmission in the APPKI model is A2AR independent.** We tested the hypothesis that the synaptic hyperexcitability at the SC terminal was driven by potentiated $A_{2A}R$ signaling. **A)** Top: Scheme of experiment to monitor SC-to-CA1 synaptic signaling (S1: stimulation; Rec: recording). Bottom: cartoon of adenosine signaling. **B)** Time trajectory of normalized fEPSP/FV before (a) and after A2AR antagonist (SCH: 50 nM). Inset: representative fEPSP traces. **C)** Mean fEPSP-F.V. relationship and **D)** individual paired slope data points (right) before (a) and after (b) dosage of the A2AR antagonist; APPKI: n= 4/4. p: p-value; d: Cohen's effect size. Mean±S.D. n=n. slices/n. mice. Statistic: generalized linear mixed effect models . Green: APPKI, Purple: APPKI+SCH.

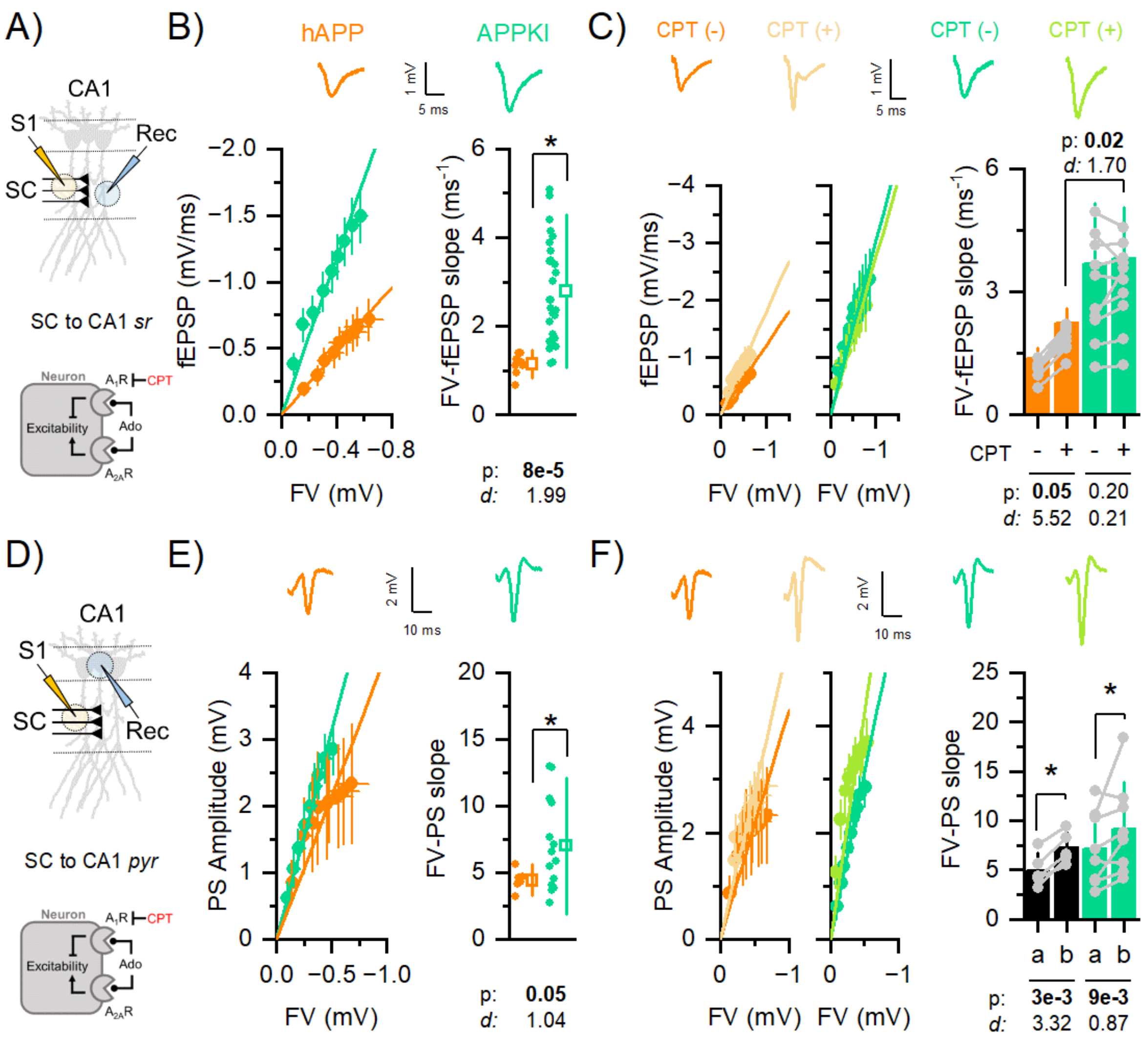


**Figure S7. Humanized sequence of APP gene does not impact adenosine signaling.** We tested the hypothesis that the humanized sequence of *APP* gene was sufficient to impact adenosine signaling. **A,D)** Top: Scheme of experiment to monitor SC-to-CA1 synaptic or somatic signaling (S1: stimulation; Rec: recording). Bottom: cartoon of adenosine signaling. **B,E)** Mean fEPSP-FV or PS-FV relationships (left) and individual slope data points (right); Synaptic; hAPP: n=10/6, APPKI: n=27/17; Somatic - hAPP: n=6/6, APPKI: n=14/10. Inset: representative fEPSP traces. **C,F)** Mean fEPSP-F.V. or PS-F.V. relationships (left) and individual paired slope data points (right) before (a) and after (b) dosage of the $A_1$R blocker; Synaptic; hAPP: n=10/6, APPKI: n=11/6; Somatic - hAPP: n=6/6, APPKI: n=8/5. The data indicated that the presence of the humanized sequence of *APP* gene was not sufficient to explain the synaptic or somatic hyperexcitability described in the APPKI model. In fact, the hAPP model was less excitable and retained basal adenosine signal. p: p-value; d: Cohen's effect size. Mean±S.D. n=n. slices/n. mice. Statistic: generalized linear mixed effect models . Orange: hAPP, Green: APPKI, Light Orange: hAPP+CPT, Light Green: APPKI+CPT.
